# Replay shapes abstract cognitive maps for efficient social navigation

**DOI:** 10.1101/2023.12.19.572418

**Authors:** Jae-Young Son, Marc-Lluís Vives, Apoorva Bhandari, Oriel FeldmanHall

## Abstract

To make adaptive social decisions, people must anticipate how information flows through their social network. While this requires knowledge of how people are connected, networks are too large to have firsthand experience with every possible route between individuals. How, then, are people able to accurately track information flow through social networks? We find that people immediately cache abstract knowledge about social network structure as they learn who is friends with whom, which enables the identification of efficient routes between remotely-connected individuals. These cognitive maps of social networks, which are built while learning, are then reshaped through overnight rest. During these extended periods of rest, a replay-like mechanism helps to make these maps increasingly abstract, which privileges improvements in social navigation accuracy for the longest communication paths that span distinct communities within the network. Together, these findings provide mechanistic insight into the sophisticated mental representations humans use for social navigation.

## MAIN

In a set of now-classic studies, Stanley Milgram asked subjects in Nebraska to forward a letter to a target individual they did not know. Subjects were only told the person’s name and that they lived in Boston. The job was to mail the letter to someone who could, in turn, forward the letter closer towards the target. Remarkably, of the letters that eventually reached their target, the source and target were only separated by about six degrees^1^. Milgram’s study illustrates the fundamental challenge of social navigation: human networks are vast yet densely-connected, meaning that a variety of things—gossip, ideas, norms, disease, and more—are susceptible to being amplified and spread by social networks^2^. To navigate this web of relationships, people must anticipate how information flows, which requires understanding how people are connected^3,4^. Although this is an inherently difficult problem, Milgram’s result suggests that people are surprisingly capable of navigating social networks, even if they lack full knowledge of how people are connected within them. Yet, despite decades of active interest, little is known about the cognitive mechanisms that enable people to solve social navigation problems.

What kinds of mental representations might support social navigation? Decades of research on spatial navigation offers a useful window into how humans might organize complex relational information. It is well-established that knowledge about physical environments is represented in cognitive maps of spatial relationships^5-7^. The format of these spatial maps allows objects to be placed within two-dimensional mental spaces^6,8^, affording representation of the longer-range relationships between those objects. Outside of spatial navigation, recent work demonstrates that humans also represent abstract maps of conceptual spaces^9-11^, including social traits such as competence and popularity^12,13^. However, relationships in social networks are poorly characterized by two-dimensional spaces, and it is not known what alternative format(s) might instead be used to build abstract cognitive maps of social networks.

Recent work in cognitive neuroscience points to a candidate representational format for social networks that encodes not only the direct connections between entities (e.g., friendships), but also longer-range, multistep connections (e.g., friends-of-friends) ^11,14-20^. This abstracted representation of social networks is related to Successor Representations in reinforcement learning, and can be learned using simple, biologically plausible mechanisms. By adjusting how many steps are integrated over, network representations can be learned at various levels of abstraction^21^, where greater abstraction confers rapid inference about distant relations, as well as the existence of network structures like communities^4,15,22,23^. This ability to represent longer-range relations likely aids social navigation, including tasks such as predicting where gossip might spread if shared with a given individual.

A second question revolves around how people efficiently build maps from limited direct experience. Evidence from rodent and human neuroscience points to an important role of replay, where the brain generalizes from experience to simulate synthetic observations that can drive additional learning, especially prioritizing those that are most critical for adaptive navigation^24-30^. Indeed, it has long been noted that a replay-like mechanism appears necessary to learn sufficiently abstract representations for navigation^15-17^. Offline replay during sleep appears to play an especially important role in generating more abstract representations of the environment^31-35^. As abstraction can help reveal the underlying structure of a given environment and therefore aid longer-range navigation^15,18^, it is likely that extended periods of rest, such as overnight sleep, are critical for building the kinds of abstract representations needed for longer-range navigation through social networks.

Therefore, multistep abstraction not only specifies a useful format for representing the topology of graph structures like social networks^4,11,16-18,23^, but also provides a natural interface between cognitive maps and replay. Although multistep abstraction is an attractive model of how people represent and navigate social networks, past research has only established that people’s memory representations of social networks are consistent with multistep abstraction^4^, and it is yet unknown whether, or how, multistep abstraction supports navigation behaviors.

Here we test whether humans rely on cognitive maps of social networks for social navigation and if a replay-like mechanism supports more successful social navigation. To assess humans’ use of cognitive maps to solve the challenge of social navigation, we created a task where subjects learn about friendships in a social network (Figure 1A), allowing us to probe how people navigate information flow through a community (Figure 1C). We then either had subjects take the navigation task immediately, or brought subjects back to the laboratory the next day to test if navigation accuracy improved after overnight rest (Figure 1F). Using computational modeling, we characterized the underlying cognitive maps employed by subjects and further tested if a replay-like mechanism helps to scaffold more successful social navigation.

**Figure 1.**
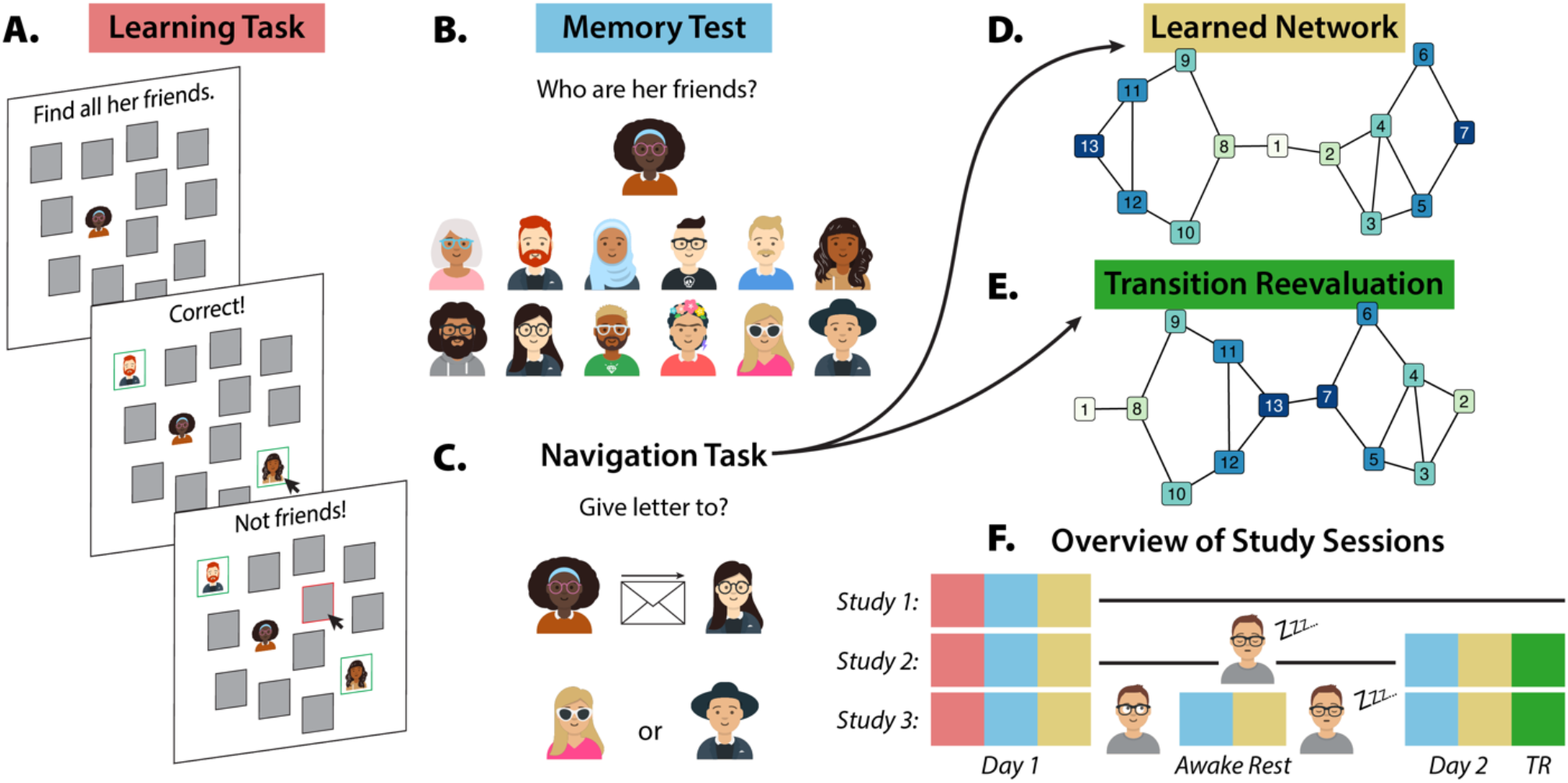
Study design. **A**. The learning task used ‘flash cards’ to facilitate rapid and accurate learning. When presented with a Target network member, subjects were required to find all of the Target’s friends from the face-down cards. When a card corresponding to the Target’s friend was selected, the card flipped face-up to reveal their photograph. Cards remained face-down in response to incorrect guesses. **B**. The memory test presented a Target and required subjects to indicate all of the Target’s friends. No feedback about accuracy was ever provided. **C**. The social navigation task presented a network member wishing to send a message to a Target through one of two Sources. Subjects were required to indicate which Source was the better choice for efficient delivery to the Target. **D**. The social network learned by subjects. **E**. In Studies 2-3, subjects were informed that some of the friendships had been broken, and that others had formed. This necessitated rapid reevaluation of how network members were related to each other. **F**. A schematic illustrating what tasks subjects completed on what days, in which studies. The color-coding corresponds to the task labels in parts A-E of this figure; yellow and green indicate completion of the navigation task for the learned and reevaluated networks, respectively. All avatar icons were generated using getavataaars.com, designed by Pablo Stanley and developed by Fang-Pen Lin.

### Humans can efficiently solve social navigation problems

We developed a novel ‘message-passing’ task as an experimental testbed of flexible social navigation, which assessed whether people understand how information flows throughout the network (Figure 1C). On each trial, a network member wished to pass a letter to a Target within the network, and needed to choose between Sources A and B. If Source A were chosen, A would pass the letter to one of their friends, who would pass it to one of *their* friends, and so on until the letter was delivered to the Target. The subject’s task was to choose the Source that would result in the most efficient delivery. Trials were classified according to the shortest path distance from the network member who had sent the letter. For example, when the correct Source was directly friends with the Target, we classified these as distance-2 problems, as the message needed to be passed twice to reach the Target. An accurate response was defined as choosing the Source with the shortest path to the Target (Methods). The Target changed from trial-to-trial, such that successful navigation required flexible use of knowledge about connections between network members. The two Sources presented on each trial were always friends of the message-sender to prevent potential confounds, and to rule out the possibility that navigation accuracy might improve simply from experience with the task, subjects were never provided with feedback.

Across three laboratory studies (total *N* = 146; data pooled for efficiency, but results replicate across all studies; Supplementary Information), subjects completed this navigation task shortly after learning friendships in a novel social network. (Figure 1A). Subjects never observed the whole network and were provided no direct information about multistep, longer-range connections between network members (e.g., friends-of-friends), but instead only observed dyadic relationships. Despite this learning format, subjects achieved above-chance navigation accuracy not only for problems where the Source was directly friends with the Target (80% accuracy at distance-2 *b* = 1.68, *Z* = 14.30, 95% CI = [1.45, 1.91], *p* < .001), but also for the longer-range problems (70% accuracy at distance-3, *b* = 1.06, *Z* = 9.34, 95% CI = [0.84, 1.28], *p* < .001; 63% accuracy at distance-4, *b* = 0.66, *Z* = 6.05, 95% CI = [0.44, 0.87], *p* < .001; Figure 2A; all results from mixed-effects logistic regression, see Methods). These results suggest that subjects learned a cognitive map that supported flexible, long-range social navigation, despite only being provided pairwise information about friendships in the network.

**Figure 2.**
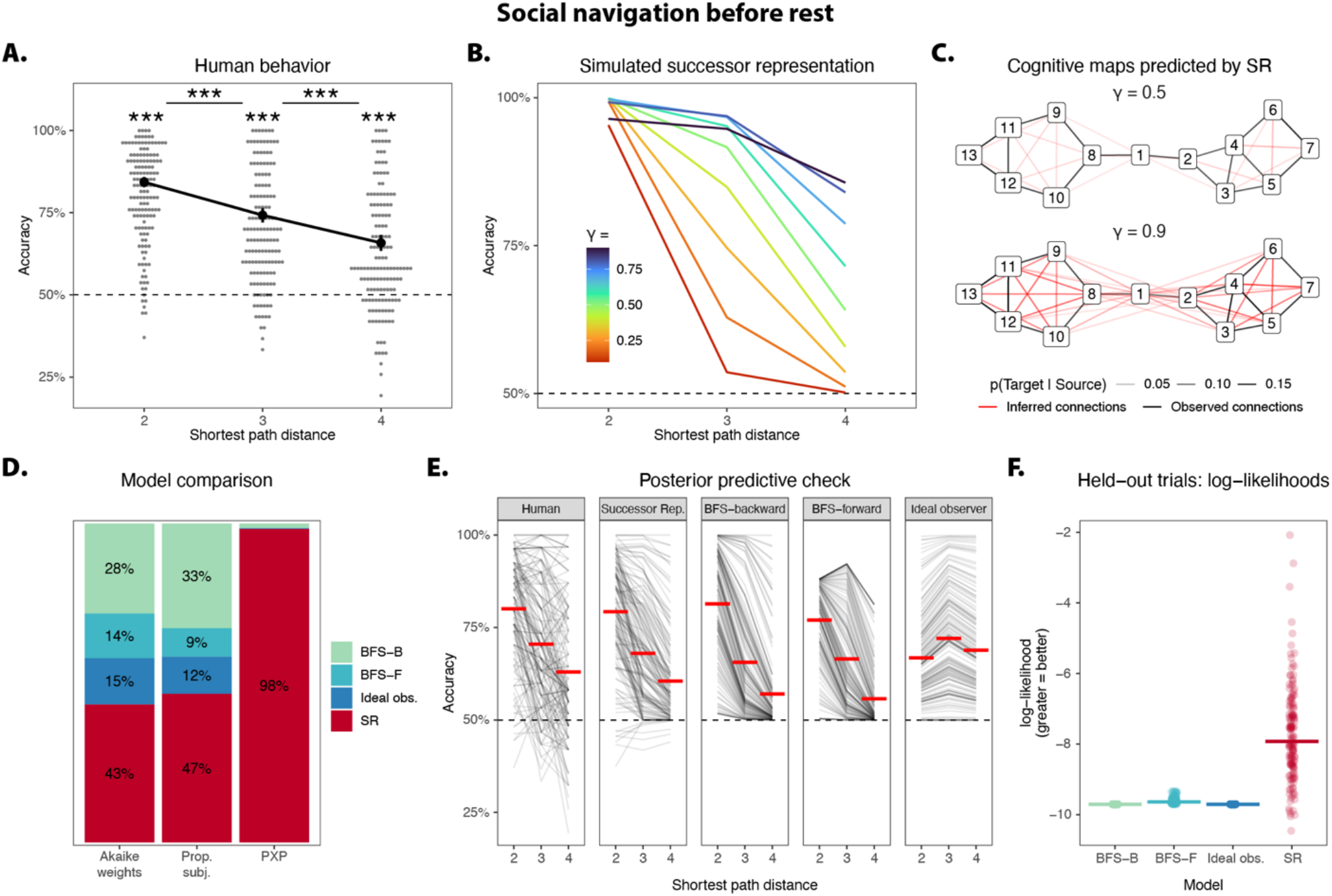
Evidence of social navigation. **A**. Shortly after learning about friendships in a novel social network, people are able to solve social navigation problems with above-chance accuracy. Trend lines reflect the estimated means from a mixed-effects logistic regression model, and error bars reflect estimated standard errors. **B**. Simulated navigation behavior from the Successor Representation (SR) model, which learns multistep relations between network members over a horizon controlled by γ. The softmax inverse temperature is fixed at β = 100 for visualization. **C**. A cognitive map learned with greater γ allows efficient representation of longer-range relationships. True friendships are color-coded in black. With increasing abstraction, the cognitive map begins to reflect inferences about community structure and connections to the ‘bridging’ nodes connecting the two communities, color-coded in red. For visualization, links have been thresholded at p(Target | Source) > .025. **D**. The SR is the best-fitting model in terms of Akaike weights, the proportion of subjects best-fit, and protected exceedance probability (PXP). **E**. Simulations of computational models’ behaviors are based on the parameters estimated from human subjects’ behaviors. Red bars reflect group-level means. **F**. In a subset of ‘held-out’ navigation problems (i.e., not used to fit parameters), subjects were presented with two Sources that were the same path distance away from the Target. Despite this, humans frequently demonstrated a preference for one Source over the other. On these held-out trials, the SR has greater out-of-sample likelihoods than the planning models, demonstrating that it is better able to explain human subjects’ preferences. Bars reflect group-level medians. **All panels**. The dashed horizontal line reflects chance-level accuracy. *** p < .001, ** p < .01, * p < .05.

### Computational models of social navigation

We consider two classes of decision-making strategies that an agent could employ to flexibly solve novel social navigation problems: model-based planning, and caching abstracted representations of multistep relations. A normatively optimal agent would represent an internal model of all pairwise friendships within the social network, then recursively iterate through those friendships until it computes the shortest path between a given Source-Target pair. In practice, online navigation of this kind is time-consuming and computationally costly^17^, but can be made tractable in small networks using search algorithms such as Breadth-First Search (BFS) that have previously been studied as cognitive ‘pathfinding’ models^36,37^. Here, we test the following planning-based models: 1) BFS-forward, in which the agent performs two forward searches from each of the two candidate Sources, choosing the Source where the Target is first found; 2) BFS-backward, in which the agent performs a single search starting from the Target, choosing the first Source that is found; and 3) an ideal observer, which computes the shortest path distance between each Source and Target, choosing the Source that is closer to the Target. To make these planning models more psychologically plausible, our implementations included parameters that captured the human tendency to give up and choose randomly during long searches, as well as decision noise when choosing between two options (Methods).

Alternatively, an agent could navigate more efficiently by caching (i.e., pre-computing) relevant knowledge. In the context of social navigation, it would be particularly useful to cache knowledge of individuals’ longer-range, multistep connections (e.g., friends-of-friends). Recent work in cognitive neuroscience points to the Successor Representation (SR) as a useful format for encoding such cached knowledge^16-18^, including cognitive maps of social networks^4^. The SR approximates the probability of transitioning from a Source to a Target in a given number of steps. A single parameter, the successor horizon γ, controls how many steps are integrated over, and therefore dictates how the SR integrates knowledge of shorter-vs longer-range connections. As γ → 0, the agent represents shorter-range relations, such that the SR only encodes one-step relations (i.e., direct friendships) when γ = 0. As γ → 1, the agent integrates over longer-range connections (e.g., friends-of-friends-of-friends…). Once the agent has cached estimates of *p*(Target | Source, γ), it can then decide between the two possible Sources using a softmax choice rule with inverse temperature β, controlling how noisily the agent chooses.

Simulation results reveal, *a priori*, that multistep abstraction is sufficient to achieve high navigation accuracy: higher values of γ were associated with greater navigation accuracy for longer-range problems, and SR agents achieved uniformly high navigation accuracy for the shorter-range problems regardless of γ (Figure 2B; Methods). These results therefore confirm that human subjects’ social navigation decisions could in principle be supported by a cognitive map of multistep relationships, where representation of longer-range relations is supported by greater abstraction (i.e., larger γ; Figure 2C).

We next fit this model to subjects’ behaviors before rest, to test whether multistep abstraction quantitatively outperforms model-based planning in explaining human behavior in the social navigation task (Methods). We used protected exceedance probability (PXP) to formally test the probability that one model provided a superior fit over all other models under consideration^38^. Results revealed that the SR indeed provided a better group-level fit to the data than all online planning models (all PXP > 0.97; Figure 2D; see Methods), mirrored in the Akaike weights^39^ and the proportion of subjects best-fit by the SR (Figure 2D). Therefore, a formal comparison of computational models suggests that human behavior on the message-passing task is best explained by the use of a cognitive map containing cached knowledge of abstract, multistep relations. Posterior predictive checks further confirmed that the planning-based models systematically mischaracterize human subjects’ navigation behaviors, while the SR model is largely successful in recapitulating human behavior (Figure 2E).

Finally, we tested whether each model was able to predict human behavior in a held-out subset of navigation problems (i.e., trials that were not used to fit model parameters) before rest. These trials were unique in that both Sources were the same shortest distance away from the Target, making them equally correct choices to a model-based agent. While the planning models do not systematically favor one Source over the other in these trials (Figure 2F), humans demonstrate preferences for Sources that have multiple (relatively) short paths to the Target, which is mirrored by the SR model predictions (Figure 2F; Supplementary Information).

### Social navigation improves with overnight rest

To test whether a replay-like mechanism might result in improved social navigation after an extended period of overnight rest that includes sleep, subjects in studies 2-3 (*N* = 96) completed a two-day procedure. The day after their first session, subjects returned to the laboratory and completed the social navigation task again. Results reveal that after overnight rest, subjects became significantly more accurate at solving problems across all distances (distance-2 accuracy 81% before rest, 82% after overnight rest, *b* = 0.23, *Z* = 2.91, 95% CI = [0.08, 0.39], *p* = .004; distance-3 accuracy 73% before rest, 75% after overnight rest, *b* = 0.26, *Z* = 2.99, 95% CI = [0.09, 0.44], *p* = .003; distance-4 accuracy 65% before rest, 71% after overnight rest, *b* = 0.43, *Z* = 5.17, 95% CI = [0.27, 0.59], *p* < .001; Figure 3A; all results from mixed-effects logistic regression). This improvement was particularly pronounced for the longest-range distance-4 problems, compared to the accuracy improvement for distance-2 problems (*b* = 0.20, *Z* = 2.36, 95% CI = [0.03, 0.36], *p* = .018).

**Figure 3.**
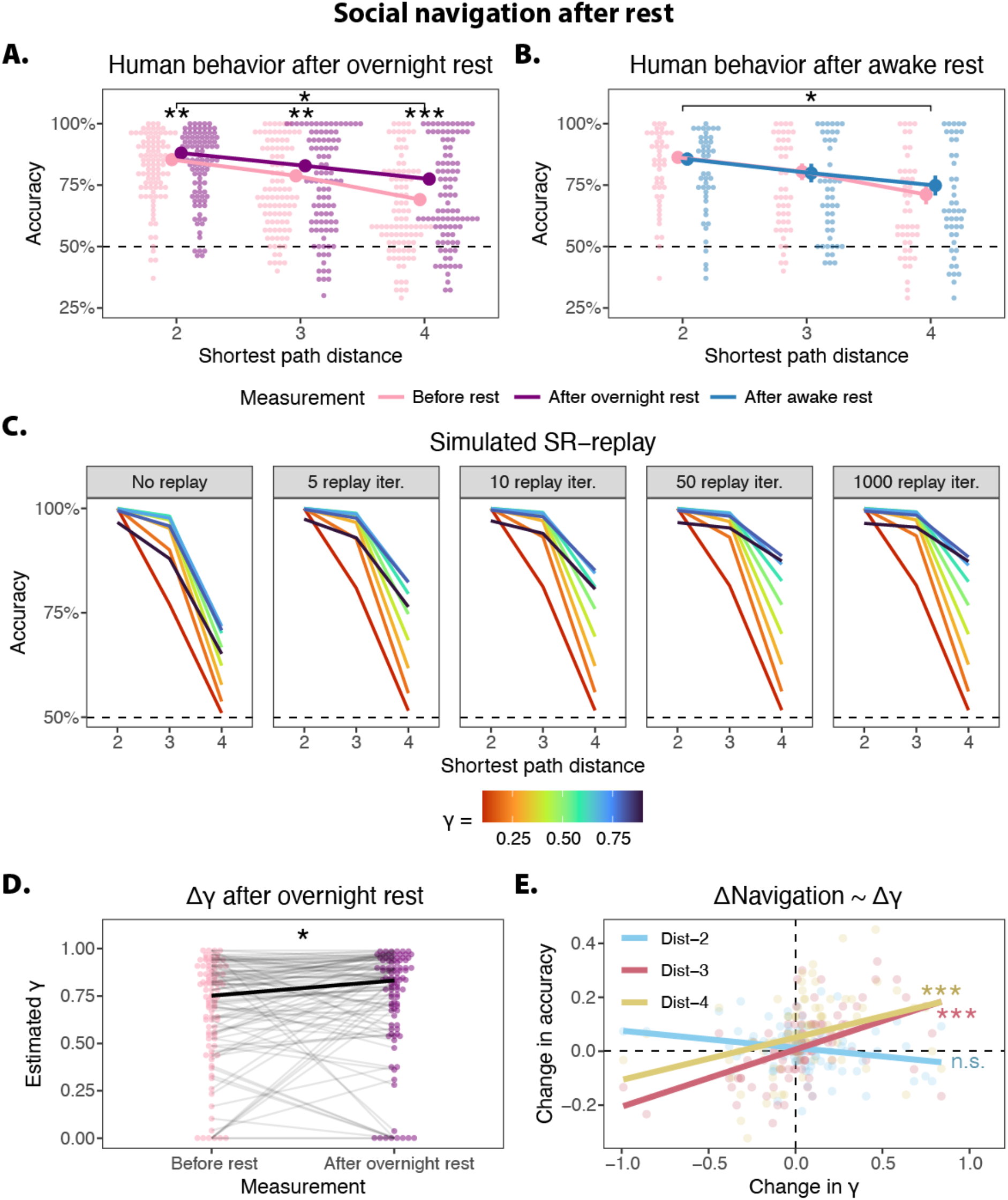
Evidence for a replay-like mechanism. **A**. After overnight rest, humans became more accurate at solving social navigation problems. Increases in navigation accuracy were particularly pronounced for problems requiring knowledge of longer-range connections. **B**. Brief awake rest did not significantly improve navigation. In panels A-B, trend lines reflect the estimated means from a mixed-effects logistic regression model, and error bars reflect estimated standard errors. **C**. Simulations suggest that even a small amount of SR-replay helps an agent consolidate its representation of friendships in a social network, resulting in more accurate navigation. However, the benefits from consolidation rapidly asymptote, and further replay does not result in notably improved navigation. The simulation results instead suggest that overnight rest may help an agent build more abstract representations (characterized by larger values of γ), which integrate over a greater number of multistep relations (e.g., friends-of-friends-of…), and which aid in solving longer-range navigation problems. **D**. Parameter estimates from the SR model fit to subjects’ navigation behaviors reveal a group-level increase in γ after overnight rest. The thick black line reflects the group-level medians. **E**. Increased γ after overnight rest was associated with improved longer-range navigation, but not shorter-range navigation. Linear trend lines are shown for visualization only; the statistical tests reflect Spearman rank-correlation. **All panels**. The dashed horizontal line reflects chance-level accuracy, except in panel E. *** p < .001, ** p < .01, * p < .05.

To test whether a brief period of awake rest is sufficient to improve navigation accuracy, subjects in study 3 (*N* = 46) were allowed to rest for approximately 15 minutes at the end of the standard Day 1 procedure and before overnight rest (Figure 1F). After this brief awake rest period, they completed the same memory and navigation tasks again. This awake rest was not sufficient for improving navigation accuracy at the group level (all Ps > .1; Figure 3B), suggesting that a longer period of rest (or possibly sleep) may be needed to produce significant improvements in navigation.

### A computational model of replay

In the theoretical framework of the Successor Representation, replay is a natural mechanism for explaining how overnight rest improves social navigation^15-17^. The knowledge cached by the SR is sensitive to an agent’s observations, which could include either direct experience from the environment or synthetic experience from offline replay. There are several plausible hypotheses of how SR-replay might result in improved navigation. For example, a ‘consolidation’ hypothesis suggests that replay fills the gaps left by insufficient direct experience, allowing the agent to learn a more stable representation^15,25^. Alternatively, an ‘abstraction’ hypothesis predicts that an agent’s ability to successfully solve longer-range navigation problems depends on building increasingly abstract representations integrating over a greater number of multistep relations (i.e., with larger γ) ^15,16,21^. Intuitively, SR-replay is likely to be less comprehensive during brief periods of awake rest, compared to extended periods of overnight rest, and it is therefore possible that overnight rest helps to stitch knowledge of pairwise relationships into representations of longer-range multistep relations, allowing an agent to build even more abstract cognitive maps.

To test these hypotheses, we conducted a simulation study examining how successfully an artificial agent could solve our social navigation problems, given varying amounts of SR-replay. For simplicity, we assumed that the agent replayed all friendships in the network in one ‘iteration’ before replaying another iteration (i.e., 17 undirected relations, 34 directed). Past empirical studies of neural replay have found that it takes approximately 50ms to replay a single transition between two states (i.e., a friendship dyad) ^24,27,40,41^, such that an SR-agent could replay 50 iterations in the span of approximately 90s. In contrast, an agent lacking an SR-replay mechanism would only be able to learn from direct experience, and would be limited to the six observations of each friendship from the learning task. We note that in translating iterations of SR-replay to absolute time, we do not imply contiguous, uninterrupted replay, but rather cumulative SR-replay that could be distributed throughout a longer period of absolute time.

Consistent with the hypothesis that SR-replay helps with consolidation, simulation results reveal that even a small amount of SR-replay is sufficient to dramatically improve navigation accuracy, compared to an agent that only learns from direct experience (Figure 3C; Methods). However, the simulation also suggests that the benefits gained from consolidation rapidly asymptote (Figure 3C). Therefore, replay-as-consolidation may help to explain how people initially achieve above-chance navigation performance, but it is unlikely to explain navigation improvement after overnight rest. Instead, consistent with an abstraction hypothesis, the simulation reveals that larger values of γ are associated with more accurate navigation decisions, especially for longer distances (Figure 2C; Figure 3C).

Given these simulation results and past research showing that a variety of mental representations become more abstract during sleep^31-35,42-46^, it may be possible that extended periods of rest enable the brain to replay longer sequences through the network during overnight rest. Empirically, this would be reflected in human navigation behaviors being better-characterized by larger values of γ after overnight rest. As hypothesized, results reveal a significant group-level increase in γ (Day 1 median γ = 0.75, Day 2 median γ = 0.83, 95% CI difference in medians [0.009, ∞], one-tailed *p* = .019; Figure 3D). To further verify that individual-level changes in estimated γ are associated with greater navigation accuracy, we used Spearman rank correlation to test whether changes in estimated γ track changes in accuracy for shorter- and longer-range navigation problems. Results reveal that increased γ on Day 2 was associated with improved navigation accuracy for the longer-range problems (distance-3 *ρ* = 0.55, one-tailed *p* < .001; distance-4 *ρ* = 0.47, one-tailed *p* < .001; Figure 3E), but not for the shorter-range problems (distance-2 *ρ* = -0.23, one-tailed *p* = .987; Figure 3E). These results are therefore consistent with the proposal that replay affords greater abstraction of a cognitive map that privileges longer-range navigation problems.

As before, we tested the alternative hypothesis that subjects’ behaviors were better-described by model-based planning. Results reveal that on both days, the SR model outperforms all planning models (all PXP > 0.97). Finally, in addition to the formal model comparison favoring the SR over the planning models, results also reveal that subjects’ memory performance significantly decreased after overnight rest (*b* = -0.29, *Z* = -4.65, 95% CI = [-0.41, -0.17], p < .001). This is inconsistent with a theory of model-based planning, as decreased memory performance suggests that an individual’s internal model gets worse, not better, following overnight rest.

### Offline gains in social navigation rely on cached structural knowledge

Why is longer-range social navigation particularly improved by building more abstract SRs? The longest navigation problems in our studies span the two communities within the network, which necessitates that information flows through an information broker connecting the groups (Figure 2C). Acquiring knowledge about possible routes passing through the broker is therefore critical for solving the longer-range navigation problems, and this knowledge becomes especially useful when information traverses across the communities. In theory, SRs built from greater values of γ should incorporate more multistep connections between the broker and other network members, thus leveraging the broker’s fundamental role in information flow. Our simulations reveal that as SRs become more abstract (with larger values of γ), information about multistep connections with the broker is cached (Figure 2C), which helps explain the observed improvement of our subjects in navigating the longest-range navigation problems. In other words, our simulations reveal *how* abstract cognitive maps extract important structural knowledge from more granular knowledge about individual friendships.

If subjects are indeed relying on cached structural knowledge to improve social navigation that spans communities, their performance should be sensitive to structural changes involving the broker, such as the broker breaking off a friendship. An agent relying on cached structural knowledge about the broker’s connections should continue to make choices as if no relationship has ruptured, since the agent will either need additional experience to learn about the network change or will need to re-cache the modified multistep relationships through replay (either offline or ‘on-task’) in order to correctly navigate the modified network. In contrast, an agent employing model-based planning would be able to rapidly incorporate that change into its internal model and alter its navigation behavior accordingly. To test whether subjects show evidence of using cached representation, we administered a transition reevaluation procedure on Day 2 in studies 2-3^16,17^ (Figure 1F). In a final task, subjects were informed that two people were no longer friends, and that two other people had newly become friends. We engineered these changes such that the critical bridge (i.e., including the broker) between the two communities was severed and formed elsewhere, while no other structural changes were made to either of the two communities (Figure 1D; Figure 1E).

Results reveal that these structural changes to the network were sufficient to abolish subjects’ improved accuracy for longer-range navigation problems after overnight rest (distance-3 *b* = 0.20, *Z* = 2.33, 95% CI = [0.03, 0.37], *p* = .020; distance-4 *b* = 0.45, *Z* = 5.51, 95% CI = [0.29, 0.61], *p* < .001), but not shorter-range problems (distance-2 *b* = 0.14, *Z* = 1.77, 95% CI = [-0.02, 0.30], *p* = .077; Figure 4A; Supplementary Information), providing evidence that subjects relied on cached structural knowledge to inform longer-range social navigation decisions. This decrease in navigation accuracy is particularly noteworthy given that subjects had, just thirty minutes prior, exhibited evidence of improved navigation following overnight rest. Indeed, after transition reevaluation, subjects’ navigation accuracy was statistically indistinguishable from their initial performance before overnight rest (Figure 4B; Supplementary Information).

**Figure 4.**
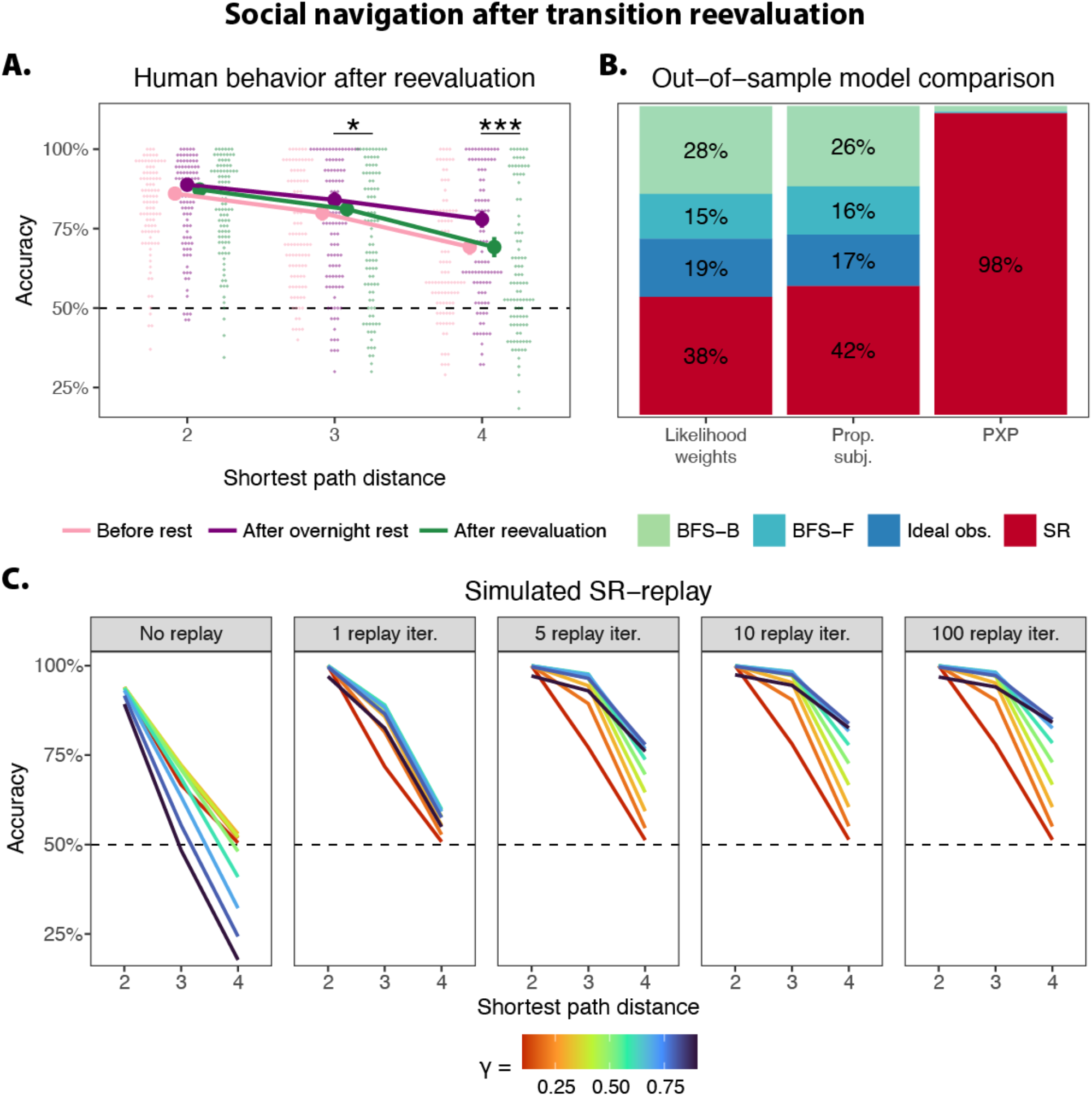
Evidence of cached structural knowledge. **A**. After being informed of changes in friendship (‘transition reevaluation’ in which the link between the bridging nodes was severed), human subjects’ navigation accuracy significantly decreased, relative to the improved navigation they had exhibited earlier that same day (i.e., after overnight rest). **B**. Using out-of-sample likelihoods to do model comparison, the SR is the best-fitting model in terms of likelihood weights (similar to Akaike weights), the proportion of subjects best-fit, and protected exceedance probability (PXP). **C**. Simulations suggest that even a small amount of on-task SR-replay helps an agent re-cache its representation of the social network after transition reevaluation, resulting in improved navigation accuracy. **All panels**. Dashed horizontal lines reflect chance-level accuracy. *** p < .001, ** p < .01, * p < .05.

To formally test whether the cached representation was more consistent with multistep abstraction, rather than model-based planning, we took advantage of the fact that subjects completed two navigation tasks on the same day (i.e., before and after transition reevaluation). This aspect of the study design enabled us to predict post-reevaluation behavior using the computational model parameters that had previously been estimated from pre-evaluation behavior, such that the data used to fit the models’ parameters was completely different from the data used to test the models’ goodness-of-fit (Methods). The SR not only has the highest out-of-sample likelihood weights (conceptually similar to Akaike weights, but based on raw log-likelihood rather than AIC), but it is also the absolute best-fitting model for the highest proportion of subjects, and is favored over all planning models in a Bayesian model selection analysis (PXP = .98; Figure 4B).

Given empirical evidence that subjects cached representations of multistep relationships, we used simulation to test how an SR-agent might use ‘on-task’ replay to quickly re-cache the structure of the network after transition reevaluation^47^. The simulation results reveal that an SR-agent that does no updating (i.e., continues to use the cached representation it had learned for the original network) is generally unable to solve distance-4 problems, with accuracy falling at, or even below, chance levels (Figure 4C). In contrast, an SR-agent that learns from replay exhibits dramatic gains in accuracy after even one iteration of replay (i.e., replaying all of the friendships in the reevaluated network; Figure 4C). Therefore, the simulation results demonstrate that even a relatively small amount of updating is sufficient for explaining how cached SRs can achieve above-chance navigation accuracy after transition reevaluation (Supplementary Information).

## Discussion

Stanley Milgram’s seminal studies, more than fifty years ago, demonstrated that people are able to efficiently pass messages through a large, complex social network, hinting at a human capacity for representing social relationships in a format that supports social navigation^1^. Here, we provide a new experimental framework for closing fundamental gaps in our mechanistic understanding of how people adaptively navigate social relationships. We find that people are proficient at solving social navigation problems requiring inference about how information spreads through a network. Indeed, people can accomplish above-chance navigation accuracy immediately after learning about a novel network, even for longer-range problems that require integrating knowledge over long chains of relationships (friends-of-friends-of-friends-of-friends). Overnight rest further improves social navigation accuracy, and has an especially pronounced effect for performance on problems involving longer-range relationships spanning different communities. Drawing inspiration from decades of research on spatial navigation in rodents and humans, we propose both a representational format enabling information flow to be tracked in the human mind, and the cognitive mechanisms for building these complex mental representations.

First, successful navigation through a network is aided by representing it as an abstract cognitive map encoding not only direct, one-step friendships, but also integrating over indirect, multistep connections like being a friend-of-a-friend. These abstract mental representations can be learned using algorithms which can extrapolate multistep relationships from disjointed, pairwise observations of friendship. People’s use of multistep abstraction allows them to build more holistic representations of how people in the network are connected to one another, and suggests that abstraction is the lynchpin of how social navigation problems are solved^3,4^.

Second, the brain further refines these cognitive maps of social networks during overnight rest using a replay-like mechanism that efficiently reuses experiences from prior learning to generate new, synthetic learning observations. This account is consistent with research showing that animals not only replay prior experiences^24,30,48^, but that they also generate entirely new, synthetic ‘walks’ through the environment^29,49^. Moreover, the fact that we observe the greatest boost in navigational improvement for long-range problems is consistent with past findings demonstrating that sleep privileges memory abstraction^31-35^. We find that after overnight rest, behavior was consistent with larger SR gammas on the second day of testing, which aligns with prior work showing that sleep appears to be important for building the kinds of highly abstract mental representations that reveal a social network’s deeper structure, such as the existence of communities and the individuals that bridge them^15^.

Our studies lay the groundwork for addressing several important questions in future work. Here, we highlight just a few of many promising directions. We establish that a replay-like mechanism is needed to explain how navigation performance improves overnight, but a fuller computational account of such a mechanism requires characterizing the content and amount of replay experienced. Past neurobiological findings strongly suggest that replay sequences consist of items that were experienced close together in time (e.g., adjacent locations in a maze that are part of the same path) ^24^. However, it remains unknown whether this holds true in the context of social networks, where an individual’s observations of social interactions may be sequentially or temporally disjointed. It also remains unknown how much neural replay is necessary for improving decision making (either in offline replay overnight, and/or on-task replay^47^), and whether a computational model could provide reasonable estimates of the amount of neural replay occurring in individual subjects. Finally, future research should test whether the benefits of SR-replay are linked to sleep specifically or can also be observed with longer periods of awake rest.

A related question revolves around the neural instantiation of multistep abstraction. Although we leverage the Successor Representation (SR) in this work, we note that multistep abstraction is a much more general representational strategy that could be implemented using many mechanisms with varying degrees of biological plausibility^9,19,20^. Despite the SR depending heavily on the temporal dynamics of experience^14,50^, multistep abstraction appears to describe how people represent social networks even when observations of social interaction are temporally disjointed^4^. It is therefore possible that the SR successfully describes social network representation because it is a useful method for discovering structure^15^, rather than being a faithful model of neural computation. A particularly intriguing possibility is that the brain may encode components of a network’s structure (i.e., basis sets) that afford greater flexibility in assembling useful representations when navigating a variety of social environments^9^. There are many ways that the brain could perform inference over graphs^20,51-53^, and it may be useful for future work to examine what kinds of basis sets are afforded by various methods of graph inference, including multistep abstraction.

In summary, people can reason about information flow efficiently in social networks by caching knowledge about long-range connections in abstracted cognitive maps. Our results provide mechanistic insights into how these abstract cognitive maps are learned, and shaped offline by a replay-like mechanism that allows the successful navigation over longer-range friendships.

## METHODS

### Subjects

In study 1, we recruited N = 50 subjects (34 female, 15 male, one nonbinary; mean age = 20.6 years old, SD = 2.81). In study 2, we recruited N = 50 subjects; one subject’s demographics were never recorded due to experimenter error. Of subjects whose demographics are known, 31 were female, 17 male, and one nonbinary; the mean age was 23.1 years old, SD = 4.63. In study 3, we recruited N = 50 subjects, but lost four datapoints due to experimenter error, leaving a final sample size of N = 46 (30 female, 16 male; mean age = 23.0 years old, SD = 4.46). All subjects received $10/hour as monetary compensation for their first study session. For the second study session, subjects in study 2 were paid $15, and subjects in study 3 were paid $20. Subjects in studies 2-3 could earn additional cash bonuses of up to $5 depending on how accurately they solved social navigation problems. All study procedures were conducted in a manner approved by the University Institutional Review Board [university name withheld for double-blind peer review], and all subjects provided informed consent.

### Overview

In study 1, a one-day study (Figure 1F), subjects first learned about a novel social network (Figure 1A), completed a memory test (Figure 1B), and then were tasked with solving social navigation problems (Figure 1C) about the network they had just learned about (Figure 1D). Details about each procedure are provided in subsequent sections.

Study 2 was a two-day study (Figure 1F), where Day 1 was identical to study 1. On Day 2, subjects returned to the lab 24 hours after their first session. In this second session, subjects completed the same memory test and social navigation task as they had in Day 1, then completed the social navigation task a third time after being informed about changes in network members’ friendships (Figure 1E).

Study 3 was a two-day study that was nearly identical to study 2, with one key modification. To test the hypothesis that brief awake rest was sufficient to improve navigation accuracy, we added a 15-minute rest period at the end of Day 1, after which subjects completed the memory test and social navigation task again (Figure 1F).

### Learning task

Subjects were required to learn the friendships within an artificial social network. To familiarize subjects with the 13 network members, the task first presented a screen presenting all network members’ faces and names, which subjects could examine for as much time as they liked. Afterwards, subjects learned about the friendships between these 13 network members from a computerized ‘flashcard’ game (Figure 1A). On each trial, subjects were shown one ‘Target’ network member, and were required to find all of the Target’s friends amongst the remaining twelve cards, which were initially displayed face-down. Subjects responded by clicking on face-down cards. Cards flipped face-up and were outlined in green when subjects made correct responses; incorrect responses were indicated by the card remaining face-down and being outlined in red (Figure 1A). Once all of the Target’s friends were identified, subjects were given three seconds to review the Target’s friends before the task moved on to the next Target.

All network members were presented as Targets, and subjects cycled through all Targets in a single block of trials before moving to the next block. The spatial mapping of network members’ cards remained consistent for the first three blocks, then was randomly shuffled for the last three blocks. This was done to ensure that subjects were truly learning about friendships and not simply spatial locations. Overall, the flashcard learning task took 20-25 minutes to complete. All face stimuli were drawn from the Chicago Face Database^54^.

### Memory test

Subjects completed a memory test immediately after the learning task (Figure 1B). Each trial presented a Target network member at the top of the screen, and all remaining network members were shown below in two rows of six photographs. Subjects responded by clicking on network members they believed to be the Target’s friend. No feedback was ever presented. All responses were self-paced, and the task took 5-10 minutes to complete.

### Social navigation task

On each trial of the ‘message-passing’ task, subjects chose between two Sources to pass a message to a given Target (Figure 1C). Subjects were explicitly informed that, depending on their choices, the message could be delivered efficiently, inefficiently, or not at all. All primary analyses were performed on trials where there was one unambiguously correct answer (based on shortest path distance). No feedback was ever presented. All responses were self-paced, and the task typically took 30-45 minutes to complete on Day 1 (Figure 1F). This procedure was identical across all three studies.

To test whether a brief period of awake rest was sufficient to improve navigation accuracy, subjects in study 3 completed the initial message-passing task on Day 1, rested for about 15 minutes, then completed the same navigation task again (Figure 1F). During the rest period, subjects completed a task that was designed to keep the social network salient in subjects’ minds, while being easy enough that subjects were actually able to rest. For the vast majority of the rest period, subjects were shown a fixation cross on a blank screen. Sporadically, the fixation cross was replaced with a photograph of a network member for 1.5 seconds. Across the entire 15-minute period, 140 photographs were displayed at random, and 50% of them were presented upside-down. The only task was to press a button when a photograph appeared upside-down.

Finally, to test for improvements in navigation performance, subjects completed the same message-passing task on Day 2 in studies 2-3 (Figure 1F). Afterwards, we administered a transition reevaluation procedure to test how alterations in network structure would impact navigation accuracy (Figure 1F). Specifically, we instructed subjects that two individuals who had previously been friends were no longer friends (Figure 1D; Figure 1E), and that a new friendship had been formed between two other network members (Figure 1E). We designed these changes specifically to break a critical bridge between two communities, and create a new bridge elsewhere. These changes in friendship invalidated the longer-range relationships cached by the SR, requiring subjects to quickly adapt to maintain high navigation accuracy. Subjects were not explicitly informed that these changes in friendship fundamentally altered the network’s structure.

### Behavioral analysis

We used the R package *glmmTMB* to estimate mixed-effects logistic regression models^55^. Whenever appropriate, we pooled data across the three studies to maximize statistical power. To account for non-independent observations, the models included random intercepts for each subject, as well as a random intercept for each study. To test memory accuracy, we pooled data from studies 2-3 and estimated a model where study session (i.e., Day 1 vs Day 2) was both a fixed-effects predictor and a random slope. In the social navigation task, shortest path distance was defined as the graph distance between the correct Source and Target after removing the ‘message-sender’ from the network (i.e., because the Source was not permitted to pass the letter back to the sender). In total, subjects completed a total of 159 trials. Of these, 14 trials presented two Sources that had the same shortest path distance (i.e., both answers were correct), 27 trials required online reevaluation of path distance (i.e., because the shortest possible path would have required sending the letter back to the sender), and 3 trials required both online reevaluation and had the same shortest path distance. Our main analyses focused on the remaining subset of 115 trials where there was an unambiguously correct answer.

We tested four behavioral hypotheses in total. Our first hypothesis tested whether subjects were able to solve navigation problems with above-chance accuracy on Day 1, immediately after learning about the social network (Figure 1F). As this procedure occurred in all three studies, we pooled data from all studies together. We included predictors for shortest path distance, which we coded as a categorical variable, and additionally estimated random slopes for shortest path distance per subject. To test whether accuracy was above chance at each distance (i.e., shortest paths of 2, 3, and 4), we iteratively re-parameterized the model by making each distance the reference category. Our second hypothesis tested whether subjects’ navigation accuracy improved on Day 2 after overnight rest (Figure 1F), and therefore pooled data from studies 2-3. This model included fixed-effects predictors for shortest path distance, the study session (i.e., Day 1 vs Day 2), and their interaction. The model also included per-subject random slopes for shortest path distance and study session. Distance-conditional changes in navigation accuracy were estimated using the same re-parameterization strategy. Our third hypothesis tested whether 15 minutes of awake rest would improve navigation accuracy. This model was functionally identical to the two-day model, except that it used data from the study 3 navigation tasks before rest and after awake rest. Finally, our fourth hypothesis tested whether changes in the network (i.e., transition reevaluation) would result in attenuated navigation accuracy using a model that was similar to the two-day model, but with two key differences: 1) the model compared navigation performance from before rest, after overnight rest, and after transition reevaluation; and 2) to control for the possible confound that the transition reevaluated trials were more difficult than the main set of navigation trials, the model included an additional predictor indexing the absolute difference in the two Sources’ shortest path distance to the Target, which serves as a proxy for task difficulty (Supplementary Information).

### Successor representation

In its typical use in reinforcement learning, the Successor Representation (SR) encodes the likelihood that an agent starting at state *s* will find itself in state *t* after taking some number of steps dictated by the successor horizon γ (i.e., the lookahead 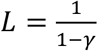). This has a straightforward translation to social navigation problems like the message-passing task, which requires computing the probability that a message given to a particular Source will be passed to the Target in some number of steps. The SR is encoded as the matrix *M* with dimensions *N* × *N*, where *N* is the number of network members. Once the SR is learned, an agent could estimate the likelihood that a message given to Source *s* will make it to Target *t* simply by looking up the value *M*(*s, t*). Here, we use function notation to index row *s* and column *t* of the matrix *M*.

In all SR-replay simulations, our implementation used a standard delta-rule method to update *M* (Equation 1; Figure 3C; Figure 4C). When network members *s* and *t* are observed together, this is encoded in the one-hot vector **1**(*t*), which is a vector of length *N* filled with zeroes except at the index *t*. The observation diverges from the agent’s prior expectation *M*(*s*) [this notation refers to the entire row *s*, as the SR retrieves and updates *M* in a row-wise manner], and therefore creates a prediction error. The agent then chains together knowledge of *s* and *t*’s friendships by adding a fractional amount of *M*(*t*) to *M*(*s*), controlled by the successor horizon γ. This overall prediction error *δ* then drives the learning update, tempered by the learning rate *α*, which was fixed to 0.1 following past work^4,16,23^. As friendships are bidirectional in our study, each learning event prompted two updates, one for *s* and another for *t*. In the simulations, we treated each novel observation as a single learning event. As the learning task consisted of six blocks, each containing learning events for both *s* → *t* and *t* → *s*, the no-replay SR learned from 12 observations of each friendship.

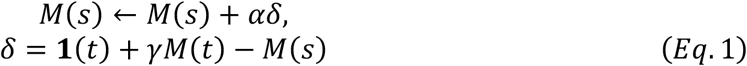

In all parameter-fitting, we used an analytic form of the SR to generate asymptotic representations (Equation 2; Figure 2B; Figure 2E), where *I* is the identity matrix, *T* is the transition matrix, and *X*^−1^ refers to the matrix inverse.

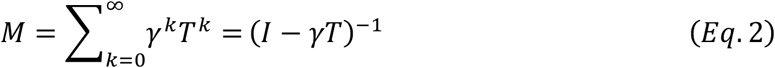

We modeled an agent’s choice between two candidate Sources using a softmax choice rule (Equation 3), such that the agent retrieves the relevant estimates from *M*(*s, t*), then probabilistically chooses the higher-valued option. Choices are made more deterministically as inverse temperature β → ∞, and more stochastically as β → 0.

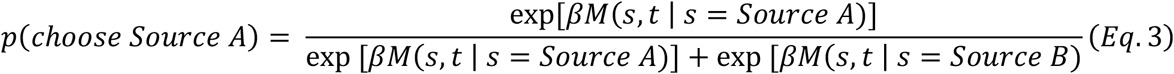

### Model-based planning

To compare multistep abstraction against model-based planning, we tested several psychologically-plausible mechanisms through which an agent could perform online planning.

The BFS-forward model (Breadth-First Search) assumes that people perform two forward searches in parallel, one from each Source, until the Target appears in one of the searches. On each iteration, the agent randomly chose to search Source A or B further. When first searching a given Source, the agent would retrieve all of that Source’s friends. Subsequent searches would retrieve friends-of-friends, friends-of-friends-of-friends, and so on. We assumed that the agent had the capacity to remember what network members had already been retrieved in each of the two searches, and the agent stopped searching once the Target was discovered. It is therefore possible that the Target may, by chance, first appear in the search associated with the incorrect answer, such that the asymptotic predictions of the BFS-forward model fall short of perfect accuracy. The BFS-forward model contains a single ‘search threshold’ parameter, which causes the agent to become increasingly likely to ‘give up’ and choose randomly when searches require iterating across long distances.

The BFS-backward model assumes that people start a single search from the Target, then iterate backward through the network until one of the two Sources is found. Backward searches are normatively more efficient than forward searches, and humans appear to prefer using backward search when it is possible for them to do so^37^. If the backward search is allowed to run to completion, this guarantees that the subject will choose the correct answer, as BFS always finds the shortest path distance between two nodes in a graph. Like BFS-forward, our implementation of BFS-backward contains a single search threshold parameter. Although the BFS-backward and -forward models use the same underlying search mechanism, we note that they make very different predictions about human behavior (Supplementary Information).

To our knowledge, there is no analytic method for computing choice and ‘reaction time’ distributions from a BFS-based search process, so we simulated how our BFS agent would solve each trial 5,000 times (description of algorithm in Supplementary Information). To fit parameters to subjects’ behavior, we defined logistic loss as the difference between a subject’s choice on a particular trial, and the average simulated choice (from 5,000 iterations) of the BFS agent on that trial (Equation 4). The search threshold parameter *τ* was estimated as the value at which a softmax with *β* = 1 was indifferent between choosing to complete a search (based on the average length of the BFS search for a given trial) and giving up (Equation 5). Likelihoods were weighted accordingly (Equation 6). For example, if an agent was estimated to be 60% likely to give up during a particularly long search, the BFS prediction contributed 40% to the overall likelihood.

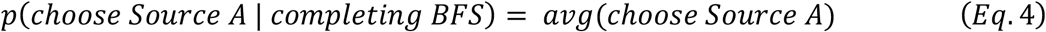

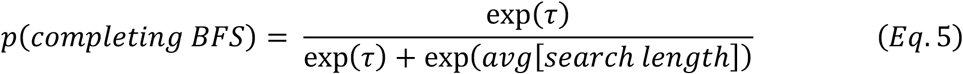

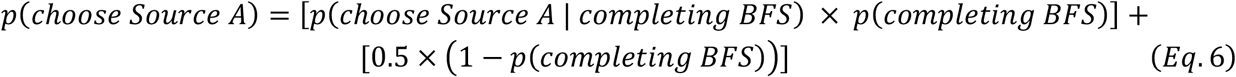

Finally, the ideal observer model assumes that people are able to compute the shortest path distances in the graph, and subsequently chooses between the two options using a softmax choice rule. We note that there are a number of psychologically-plausible processes an agent could use to compute shortest path distances, including backward-BFS. Our goal is not to adjudicate between different process models, but rather to test whether any such ideal observer could provide a compelling alternative explanation for our empirical results. This model contains a single parameter, the softmax inverse temperature *β*, which controls the agent’s sensitivity to the Sources’ shortest path distances from the Target (Equation 7).

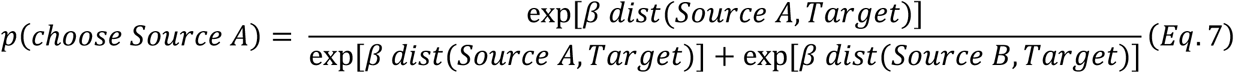

### Computational modeling

Maximum-likelihood parameter-fitting was performed using R’s default optimizer, using the ‘Nelder-Mead’ algorithm for the two-parameter SR, and the ‘Brent’ algorithm for all the single-parameter planning models. Parameters were fit independently for each subject. The SR model was re-estimated 25 times, keeping only the estimates that best maximized the likelihood; single-parameter models were only estimated once, as the fitting procedure is akin to grid search. Plots of all estimated parameters can be found in the Supplementary Information.

To verify that parameter estimates are psychologically meaningful, we performed parameter recovery and model confusability analyses for all models under consideration (Supplementary Information) ^56^. Parameter recovery analyses indicate that all key parameters are straightforwardly interpretable, and the model confusability analyses confirm that the parameter-fitting procedure is not biased in favor of our hypothesis.

In our primary model comparison procedure, we found converging evidence from three metrics: Akaike weights, the proportion of subjects best-fit by a particular model, and protected exceedance probabilities (PXP). Two of these metrics are more descriptive: Akaike weights quantify the conditional probabilities for each model based on differences in AIC^39^, and the proportion of subjects best-fit by a particular model provides an intuitive sense for models’ goodness-of-fit. Protected exceedance probabilities (PXP) provide a formal test of a model’s group-level fit compared to other candidate models^38^, and were computed using R software written by Matteo Lisi (https://github.com/mattelisi/bmsR). As our measure of log-evidence, we used the Akaike Information Criterion corrected for relatively few datapoints (i.e., AICc), which penalizes models in proportion to the number of free parameters estimated^39^.

Using the parameters estimated from the primary trials of interest (detailed in the section on behavioral analysis), we tested how well each model is explain to explain held-out data by computing out-of-sample likelihoods. This was done by simulating the models’ predictions given the estimated parameters, then computing the resulting logistic loss based on subjects’ actual behaviors in the held-out data. After computing out-of-sample likelihoods for the transition reevaluated navigation trials, we performed formal model comparison as detailed above, except using log-likelihoods rather than AICc, as no parameters were fit for out-of-sample prediction.

## ACKNOWLEDGEMENTS

We thank the following people for assisting with data collection: Isabella Aslarus, Kayleigh Danowski, Elizabeth Duchan, Yi-Fei Jerry Hu, Alexus Lawrence, Jonathan Palfy, Vera Poyraz, Mehak Malhotra, Maya Mazumder, Samantha Shulman, Ariel Stein, Sofía Vaca Narvaja, and Jenny Wang. We thank Armin Maddah for developing some of the task code used in these studies, and Yi Yang Teoh for helpful computational modeling advice. Part of this research was conducted using computational resources and services at the Center for Computation and Visualization, Brown University. Advanced access to these computing resources was supported by NIH award 1S10OD025181. This work is supported by the National Science Foundation award 2123469 (O.F.H. and A.B.).

## DATA AND CODE AVAILABILITY

All data and code needed to reproduce the analyses are available in a publicly-accessible GitHub repository: https://github.com/feldmanhalllab/network-navigation-replay

## AUTHOR CONTRIBUTIONS

Conceptualization: M-L.V., J.Y.S., A.B., O.F.H. Formal analysis: J.Y.S., M-L.V. Funding acquisition: O.F.H. Investigation: M-L.V. Methodology: M-L.V., J.Y.S., A.B., O.F.H. Supervision: O.F.H., A.B., Writing: J.Y.S., M-L.V., A.B., O.F.H.

## REFERENCES

1 Traver, J. & Milgram, S. An Experimental Study of the Small World Problem. Sociometry 32, 425–443 (1969). 10.2307/2786545

2 Christakis, N. A. & Fowler, J. H. Social contagion theory: examining dynamic social networks and human behavior. Statistics in Medicine 32, 556–577 (2013). 10.1002/sim.5408

3 Son, J.-Y., Bhandari, A. & FeldmanHall, O. Cognitive maps of social features enable flexible inference in social networks. Proceedings of the National Academy of Sciences 118, e2021699118 (2021). 10.1073/pnas.2021699118

4 Son, J.-Y., Bhandari, A. & FeldmanHall, O. Abstract cognitive maps of social network structure aid adaptive inference. Proceedings of the National Academy of Sciences 120, e2310801120 (2023). 10.1073/pnas.2310801120

5 Tolman, E. C. Cognitive maps in rats and men. Psychological Review 55, 189–208 (1948). 10.1037/h0061626

6 O’Keefe, J. & Nadel, L. The Hippocampus as a Cognitive Map. (Oxford: Clarendon Press, 1978).

7 Hafting, T., Fyhn, M., Molden, S.Moser, M.-B. & Moser, E. I. Microstructure of a spatial map in the entorhinal cortex. Nature 436, 801–806 (2005). 10.1038/nature03721

8 Bellmund, J. L. S., Gärdenfors, P., Moser, E. I. & Doeller, C. F. Navigating cognition: Spatial codes for human thinking. Science 362, eaat6766 (2018). 10.1126/science.aat6766

9 Behrens, T. E. J. et al. What Is a Cognitive Map? Organizing Knowledge for Flexible Behavior. Neuron 100, 490–509 (2018). 10.1016/j.neuron.2018.10.002

10 Constantinescu, A. O., O’Reilly, J. X. & Behrens, T. E. J. Organizing conceptual knowledge in humans with a gridlike code. Science 352, 1464 (2016). 10.1126/science.aaf0941

11 Garvert, M. M., Dolan, R. J. & Behrens, T. E. J. A map of abstract relational knowledge in the human hippocampal–entorhinal cortex. eLife 6, e17086 (2017). 10.7554/eLife.17086

12 Tavares, Rita M. et al. A Map for Social Navigation in the Human Brain. Neuron 87, 231–243 (2015). 10.1016/j.neuron.2015.06.011

13 Park, S. A., Miller, D. S., Nili, H., Ranganath, C. & Boorman, E. D. Map Making: Constructing, Combining, and Inferring on Abstract Cognitive Maps. Neuron 107, 1226-1238.e1228 (2020). 10.1016/j.neuron.2020.06.030

14 Dayan, P. Improving Generalization for Temporal Difference Learning: The Successor Representation. Neural Computation 5, 613–624 (1993). 10.1162/neco.1993.5.4.613

15 Momennejad, I. Learning Structures: Predictive Representations, Replay, and Generalization. Current Opinion in Behavioral Sciences 32, 155–166 (2020). 10.1016/j.cobeha.2020.02.017

16 Momennejad, I. et al. The successor representation in human reinforcement learning. Nature Human Behaviour 1, 680–692 (2017). 10.1038/s41562-017-0180-8

17 Russek, E. M., Momennejad, I., Botvinick, M. M., Gershman, S. J. & Daw, N. D. Predictive representations can link model-based reinforcement learning to model-free mechanisms. PLOS Computational Biology 13, e1005768 (2017). 10.1371/journal.pcbi.1005768

18 Stachenfeld, K. L., Botvinick, M. M. & Gershman, S. J. The hippocampus as a predictive map. Nature Neuroscience 20, 1643–1653 (2017). 10.1038/nn.4650

19 Lynn, C. W. & Bassett, D. S. How humans learn and represent networks. Proceedings of the National Academy of Sciences 117, 29407 (2020). 10.1073/pnas.1912328117

20 Lynn, C. W., Kahn, A. E., Nyema, N. & Bassett, D. S. Abstract representations of events arise from mental errors in learning and memory. Nature Communications 11, 2313 (2020). 10.1038/s41467-020-15146-7

21 Momennejad, I. & Howard, M. W. Predicting the Future with Multi-scale Successor Representations. bioRxiv, 449470 (2018). 10.1101/449470

22 Schapiro, A. C., Rogers, T. T., Cordova, N. I., Turk-Browne, N. B. & Botvinick, M. M. Neural representations of events arise from temporal community structure. Nature Neuroscience 16, 486–492 (2013). 10.1038/nn.3331

23 Pudhiyidath, A. et al. Representations of Temporal Community Structure in Hippocampus and Precuneus Predict Inductive Reasoning Decisions. Journal of Cognitive Neuroscience, 1–25 (2022). 10.1162/jocn_a_01864

24 Foster, D. J. Replay Comes of Age. Annual Review of Neuroscience 40, 581–602 (2017). 10.1146/annurev-neuro-072116-031538

25 Schapiro, A. C., McDevitt, E. A., Rogers, T. T., Mednick, S. C. & Norman, K. A. Human hippocampal replay during rest prioritizes weakly learned information and predicts memory performance. Nature Communications 9, 3920 (2018). 10.1038/s41467-018-06213-1

26 Sun, W., Advani, M., Spruston, N., Saxe, A. & Fitzgerald, J. E. Organizing memories for generalization in complementary learning systems. Nature Neuroscience 26, 1438–1448 (2023). 10.1038/s41593-023-01382-9

27 Liu, Y., Mattar, M. G., Behrens, T. E. J., Daw, N. D. & Dolan, R. J. Experience replay is associated with efficient nonlocal learning. Science 372, eabf1357 (2021). 10.1126/science.abf1357

28 Foster, D. J. & Wilson, M. A. Reverse replay of behavioural sequences in hippocampal place cells during the awake state. Nature 440, 680–683 (2006). 10.1038/nature04587

29 Igata, H., Ikegaya, Y. & Sasaki, T. Prioritized experience replays on a hippocampal predictive map for learning. Proceedings of the National Academy of Sciences 118, e2011266118 (2021). 10.1073/pnas.2011266118

30 Zhenglong, Z., Michael, J. K. & Anna, C. S. Replay as context-driven memory reactivation. bioRxiv, 2023.2003.2022.533833 (2023). 10.1101/2023.03.22.533833

31 Ellenbogen, J. M., Hu, P. T., Payne, J. D., Titone, D. & Walker, M. P. Human relational memory requires time and sleep. Proceedings of the National Academy of Sciences 104, 7723–7728 (2007). 10.1073/pnas.0700094104

32 Lewis, P. A. & Durrant, S. J. Overlapping memory replay during sleep builds cognitive schemata. Trends in Cognitive Sciences 15, 343–351 (2011). 10.1016/j.tics.2011.06.004

33 Lutz, N. D., Diekelmann, S., Hinse-Stern, P., Born, J. & Rauss, K. Sleep Supports the Slow Abstraction of Gist from Visual Perceptual Memories. Scientific Reports 7, 42950 (2017). 10.1038/srep42950

34 Feld, G. B., Bernard, M., Rawson, A. B. & Spiers, H. J. Sleep targets highly connected global and local nodes to aid consolidation of learned graph networks. Scientific Reports 12, 15086 (2022). 10.1038/s41598-022-17747-2

35 Klinzing, J. G., Niethard, N. & Born, J. Mechanisms of systems memory consolidation during sleep. Nature Neuroscience 22, 1598–1610 (2019). 10.1038/s41593-019-0467-3

36 Correa, C. G., Ho, M. K., Callaway, F., Daw, N. D. & Griffiths, T. L. Humans decompose tasks by trading off utility and computational cost. PLOS Computational Biology 19, e1011087 (2023). 10.1371/journal.pcbi.1011087

37 Callaway, F. et al. Rational use of cognitive resources in human planning. Nature Human Behaviour 6, 1112–1125 (2022). 10.1038/s41562-022-01332-8

38 Rigoux, L., Stephan, K. E., Friston, K. J. & Daunizeau, J. Bayesian model selection for group studies — Revisited. NeuroImage 84, 971–985 (2014). 10.1016/j.neuroimage.2013.08.065

39 Wagenmakers, E.-J. & Farrell, S. AIC model selection using Akaike weights. Psychonomic Bulletin & Review 11, 192–196 (2004). 10.3758/BF03206482

40 Kurth-Nelson, Z., Economides, M., Dolan Raymond J. & Dayan, P. Fast Sequences of Non-spatial State Representations in Humans. Neuron 91, 194–204 (2016). 10.1016/j.neuron.2016.05.028

41 Schuck, N. W. & Niv, Y. Sequential replay of nonspatial task states in the human hippocampus. Science 364, eaaw5181 (2019). 10.1126/science.aaw5181

42 Gómez, R. L., Bootzin, R. R. & Nadel, L. Naps Promote Abstraction in Language-Learning Infants. Psychological Science 17, 670–674 (2006). 10.1111/j.1467-9280.2006.01764.x

43 Lau, H., Alger, S. E. & Fishbein, W. Relational Memory: A Daytime Nap Facilitates the Abstraction of General Concepts. PLOS ONE 6, e27139 (2011). 10.1371/journal.pone.0027139

44 Pereira, S. I. R. et al. Rule Abstraction Is Facilitated by Auditory Cuing in REM Sleep. The Journal of Neuroscience 43, 3838 (2023). 10.1523/JNEUROSCI.1966-21.2022

45 St Clair, M. C. & Monaghan, P. 30 edn.

46 Walker, M. P. & Stickgold, R. Overnight alchemy: sleep-dependent memory evolution. Nature Reviews Neuroscience 11, 218–218 (2010). 10.1038/nrn2762-c1

47 Wittkuhn, L., Krippner, L. M. & Schuck, N. W., Statistical learning of successor representations is related to on-task replay. bioRxiv, 2022.2002.2002.478787 (2022). 10.1101/2022.02.02.478787

48 Jadhav, S. P., Kemere, C., German, P. W. & Frank, L. M. Awake Hippocampal Sharp-Wave Ripples Support Spatial Memory. Science 336, 1454–1458 (2012). 10.1126/science.1217230

49 Stoianov, I., Maisto, D. & Pezzulo, G. The hippocampal formation as a hierarchical generative model supporting generative replay and continual learning. Progress in Neurobiology 217, 102329 (2022). 10.1016/j.pneurobio.2022.102329

50 Gershman, S. J., Moore, C. D., Todd, M. T., Norman, K. A. & Sederberg, P. B. The Successor Representation and Temporal Context. Neural Computation 24, 1553–1568 (2012). 10.1162/NECO_a_00282

51 Lau, T., Gershman, S. J. & Cikara, M. Social structure learning in human anterior insula. eLife 9, e53162 (2020). 10.7554/eLife.53162

52 Whittington, J. C. R. et al. The Tolman-Eichenbaum Machine: Unifying Space and Relational Memory through Generalization in the Hippocampal Formation. Cell 183, 1249-1263.e1223 (2020). 10.1016/j.cell.2020.10.024

53 Wu, C. M., Schulz, E. & Gershman, S. J. Inference and Search on Graph-Structured Spaces. Computational Brain & Behavior 4, 125–147 (2021). 10.1007/s42113-020-00091-x

54 Ma, D. S., Correll, J. & Wittenbrink, B. The Chicago face database: A free stimulus set of faces and norming data. Behavior Research Methods 47, 1122–1135 (2015). 10.3758/s13428-014-0532-5

55 Brooks, M. E. et al. glmmTMB balances speed and flexibility among packages for zero-inflated generalized linear mixed modeling. The R journal 9, 378–400 (2017).

56 Wilson, R. C. & Collins, A. G. E. Ten simple rules for the computational modeling of behavioral data. eLife 8, e49547 (2019). 10.7554/eLife.49547

