## Supplementary Information for "Replay shapes abstract cognitive maps for efficient social navigation"

### BEHAVIORAL ANALYSES

*Memory accuracy before and after overnight rest.*

Disaggregated results from studies 1-3 are available in Table S1.

| *Study* | *Term* | *Beta estimate* | *Z* | *95% CI* | *p* |
|---|---|---|---|---|---|
| 1 | Intercept | 2.36 | 16.06 | [2.08, 2.65] | < .001 |
| 2 | Intercept (BOR) | 2.80 | 17.44 | [2.48, 3.11] | < .001 |
| | AOR | -0.25 | -4.29 | [-0.36, -0.13] | < .001 |
| 3 | Intercept (BOR) | 2.77 | 12.87 | [2.35, 3.19] | < .001 |
| | AOR | -0.21 | -3.76 | [-0.32, -0.10] | < .001 |

***Table S1 | Memory accuracy before and after overnight rest.*** *A mixed-effects logistic regression model was estimated separately for each study. All models included random intercepts. 'Before overnight rest' is abbreviated BOR, and 'after overnight rest' is abbreviated AOR.*

*Memory accuracy after awake rest.*

In study 3, subjects completed a memory test after awake rest (Figure 1F). A statistical comparison of memory accuracy after awake rest, before overnight rest, and after overnight rest is provided in Table S2.

| *Study* | *Term* | *Beta estimate* | *Z* | *95% CI* | *p* |
|---|---|---|---|---|---|
| 3 | Intercept (AAR) | 2.64 | 11.65 | [2.20, 3.08] | < .001 |
| | BOR | 0.17 | 2.94 | [0.06, 0.28] | .003 |
| | AOR | -0.05 | -0.82 | [-0.15, 0.06] | .411 |

***Table S2 | Memory accuracy after awake rest.*** *Results from mixed-effects logistic regression model including random intercepts. 'After awake rest' is abbreviated AAR, 'before overnight rest' is abbreviated BOR, and 'after overnight rest' is abbreviated AOR.*

*Social navigation before overnight rest.*

In our analyses of social navigation behaviors, we treated shortest path distance as a categorical variable. Therefore, to test whether subjects achieved above-chance navigation accuracy at each distance, we iteratively re-parameterized the regression model so that the reference category reflected estimates for each distance. As a consequence, many of the estimates are redundant with each other. For the sake of efficiency, we report all disaggregated, non-redundant estimates in Table S3.

| ***Study*** | ***Reference category*** | ***Term*** | ***Beta estimate*** | ***Z*** | ***95% CI*** | ***p*** |
|---|---|---|---|---|---|---|
| 1 | Distance 2 | Intercept | 1.34 | 12.76 | [1.14, 1.55] | < .001 |
| | | Distance 3 | -0.66 | -8.90 | [-0.80, -0.51] | < .001 |
| | | Distance 4 | -0.93 | -12.92 | [-1.07, -0.79] | < .001 |
| | Distance 3 | Intercept | 0.69 | 6.29 | [0.47, 0.90] | < .001 |
| | | Distance 4 | -0.28 | -3.53 | [-0.43, -0.12] | < .001 |
| | Distance 4 | Intercept | 0.41 | 3.82 | [0.20, 0.62] | < .001 |
| 2 | Distance 2 | Intercept | 1.71 | 12.77 | [1.45, 1.98] | < .001 |
| | | Distance 3 | -0.57 | -7.17 | [-0.73, -0.42] | < .001 |
| | | Distance 4 | -1.01 | -13.15 | [-1.16, -0.86] | < .001 |
| | Distance 3 | Intercept | 1.14 | 8.28 | [0.87, 1.41] | < .001 |
| | | Distance 4 | -0.44 | -5.26 | [-0.60, -0.27] | < .001 |
| | Distance 4 | Intercept | 0.70 | 5.19 | [0.44, 0.97] | < .001 |
| 3 | Distance 2 | Intercept | 1.81 | 9.76 | [1.45, 2.18] | < .001 |
| | | Distance 3 | -0.46 | -5.37 | [-0.63, -0.29] | < .001 |
| | | Distance 4 | -0.92 | -11.21 | [-1.08, -0.76] | < .001 |
| | Distance 3 | Intercept | 1.36 | 7.16 | [0.98, 1.73] | < .001 |
| | | Distance 4 | -0.46 | -5.13 | [-0.64, -0.29] | < .001 |
| | Distance 4 | Intercept | 0.89 | 4.77 | [0.53, 1.26] | < .001 |

***Table S3 | Social navigation before overnight rest.*** *Results from mixed-effects logistic regression models including random intercepts.*

*Social navigation after overnight rest.*

We took a similar approach to analyzing social navigation after overnight rest for studies 2-3; disaggregated, non-redundant results are reported in Table S4.

| *Study* | *Reference category* | *Term* | *Beta estimate* | *Z* | *95% CI* | *p* |
|---|---|---|---|---|---|---|
| 2 | Distance 2 | Intercept (BOR) | 1.72 | 12.68 | [1.45, 1.98] | < .001 |
| | | Distance 3 | -0.57 | -7.17 | [-0.73, -0.42] | < .001 |
| | | Distance 4 | -1.01 | -13.15 | [-1.16, -0.86] | < .001 |
| | | AOR | 0.19 | 1.86 | [-0.01, 0.40] | .063 |
| | | Distance 3 × AOR | 0.13 | 1.13 | [-0.10, 0.36] | .258 |
| | | Distance 4 × AOR | 0.24 | 2.14 | [0.02, 0.46] | .032 |
| | Distance 3 | Intercept | 1.14 | 8.23 | [0.87, 1.41] | < .001 |
| | | Distance 4 | -0.44 | -5.26 | [-0.60, -0.27] | < .001 |
| | | AOR | 0.33 | 2.86 | [0.10, 0.55] | .004 |
| | | Distance 4 × AOR | 0.11 | 0.89 | [-0.13, 0.35] | .376 |
| | Distance 4 | Intercept | 0.70 | 5.17 | [0.44, 0.97] | < .001 |
| | | AOR | 0.43 | 4.02 | [0.22, 0.64] | < .001 |
| 3 | Distance 2 | Intercept (BOR) | 1.82 | 9.60 | [1.45, 2.20] | < .001 |
| | | Distance 3 | -0.46 | -5.36 | [-0.63, -0.29] | < .001 |
| | | Distance 4 | -0.92 | -11.21 | [-1.08, -0.76] | < .001 |
| | | AOR | 0.29 | 2.37 | [0.05, 0.54] | .018 |
| | | Distance 3 × AOR | -0.15 | -1.19 | [-0.39, 0.10] | .235 |
| | | Distance 4 × AOR | 0.16 | 1.32 | [-0.08, 0.39] | .187 |
| | Distance 3 | Intercept | 1.37 | 7.06 | [0.99, 1.74] | < .001 |
| | | Distance 4 | -0.46 | -5.13 | [-0.64, -0.29] | < .001 |
| | | AOR | 0.15 | 1.11 | [-0.11, 0.41] | .267 |
| | | Distance 4 × AOR | 0.30 | 2.33 | [0.05, 0.56] | .020 |
| | Distance 4 | Intercept | 0.90 | 4.73 | [0.53, 1.28] | < .001 |
| | | AOR | 0.45 | 3.53 | [0.20, 0.70] | < .001 |

***Table S4 | Social navigation after overnight rest.*** *Results from mixed-effects logistic regression models including random intercepts and random slopes for the measurement ID (i.e., before overnight rest, after overnight rest). 'Before overnight rest' is abbreviated BOR, and 'after overnight rest' is abbreviated AOR. Note that the reference category was set to BOR, such that all 'main effects' reflect estimates for BOR, and all 'interaction' effects reflect estimates for AOR.*

*Social navigation after awake rest.*

Only subjects in study 3 completed the social navigation task after awake rest. The full table of results is reported in Table S5.

| *Study* | *Reference category* | *Term* | *Beta estimate* | *Z* | *95% CI* | *p* |
|---|---|---|---|---|---|---|
| 3 | Distance 2 | Intercept (BOR) | 1.84 | 9.54 | [1.46, 2.22] | < .001 |
| | | Distance 3 | -0.42 | -3.29 | [-0.68, -0.17] | .001 |
| | | Distance 4 | -0.94 | -6.97 | [-1.20, -0.67] | < .001 |
| | | AAR | -0.06 | -0.48 | [-0.30, 0.18] | .635 |
| | | Distance 3 × AAR | 0.02 | 0.18 | [-0.22, 0.26] | .861 |
| | | Distance 4 × AAR | 0.24 | 2.06 | [0.01, 0.48] | .039 |
| | Distance 3 | Intercept | 1.42 | 6.71 | [1.00, 1.83] | < .001 |
| | | Distance 4 | -0.51 | -4.43 | [-0.74, -0.29] | < .001 |
| | | AAR | -0.04 | -0.27 | [-0.30, 0.22] | .785 |
| | | Distance 4 × AAR | 0.22 | 1.72 | [-0.03, 0.48] | .085 |
| | Distance 4 | Intercept | 0.90 | 4.68 | [0.53, 1.28] | < .001 |
| | | AAR | 0.19 | 1.48 | [-0.06, 0.43] | .140 |

***Table S5 | Social navigation after awake rest.*** *Results from mixed-effects logistic regression models including random intercepts and random slopes for measurement ID (i.e., before overnight rest, after awake rest), as well as distance. 'Before overnight rest' is abbreviated BOR, and 'after awake rest' is abbreviated AAR. Note that the reference category was set to BOR, such that all 'main effects' reflect estimates for BOR, and all 'interaction' effects reflect estimates for AAR.*

### COMPUTATIONAL MODELS

*Breadth-First Search (BFS).*

The core algorithm used to perform BFS was adapted from standard implementations that are common in cognitive and computer science. In a single 'iteration' of BFS, the agent performs the following steps:

1. Retrieve the first element (i.e., node) in the search queue, and search for its neighbors (i.e., nodes that it is directly connected to)
2. Identify the neighbors who have not yet been visited, and add them to the search queue in a random order
3. Mark the neighbors as having been visited
4. De-queue the first element from the queue (i.e., the node that had just been searched)

If the Target has not been found (i.e., visited), and the queue is non-empty, the search continues with another iteration of BFS. In our simulations, we recorded information about the number of visits needed to fully complete a search (i.e., the 'search length'). As detailed in the Methods, we then estimated a 'search threshold' parameter for the BFS-agents, which allows the agent to probabilistically choose between continuing a search (if its threshold is greater than the search length), versus giving up and choosing between the two Sources randomly.

*A priori computational model simulations.*

The set of candidate models was chosen for their psychological plausibility, as well their ability to provide insights into possible cognitive mechanisms underlying social navigation. Though the posterior predictive check provides an intuitive visualization of these models' best-fitting predictions of human behavior, visualizations of *a priori* model simulations provide an intuition for how different parameter settings influence predicted the planning models' behaviors (Figure S1).

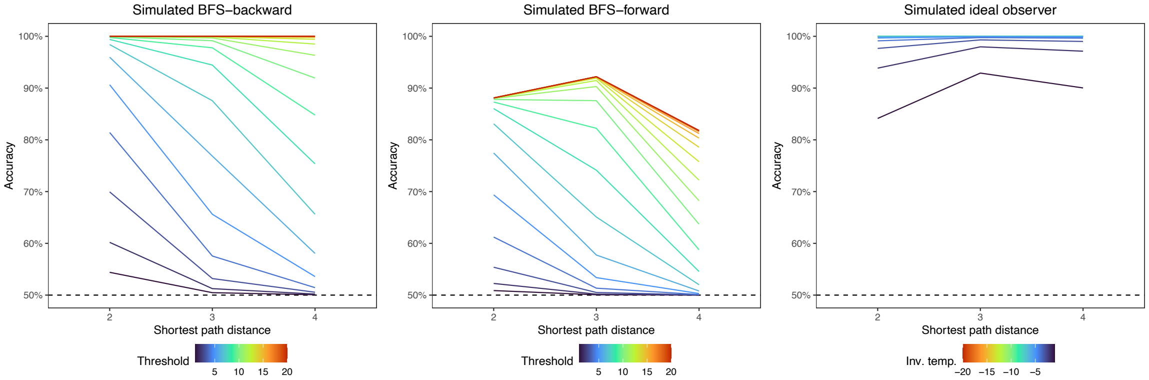


***Figure S1 | A priori simulations of planning models.*** *Parameter values were selected based on when each model's predictions became saturated (i.e., reached asymptote).*

There are two different methods of implementing of the SR, which either compute an asymptotic SR using an analytic, closed-form solution, or use a delta-rule updating mechanism to build an SR from observation. *A priori* simulations indicate that these implementations can in principle make different predictions (Figure S2), particularly at smaller values of $\gamma$.

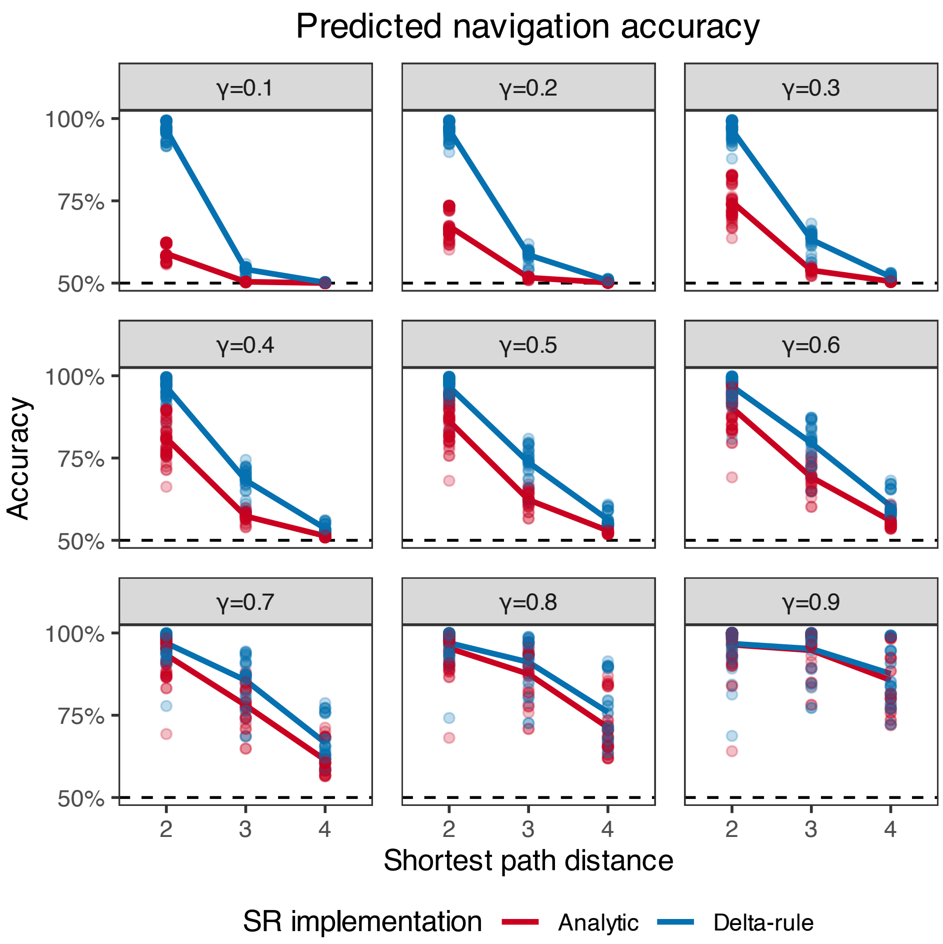


***Figure S2 | A priori simulations of analytic and delta-rule SRs.*** *The model predictions are most divergent for smaller values of* $\gamma$*, and converge as* $\gamma$ *becomes larger. Each datapoint reflects one trial from the navigation task.*

### PARAMETER-FITTING

*Parameter recovery.*

Before performing the parameter recovery analysis, we simulated behavior from all computational models at various parameter values to test what parameter ranges produced meaningful changes in predicted behavior. We found that the BFS-backward model saturated at a search threshold of 15 (i.e., produced asymptotic behavior, after which larger search thresholds did not make a difference), that the BFS-forward model saturated at a search threshold of 16, that the ideal observer saturated at a softmax inverse temperature of -5 (i.e., produced meaningful behavior when the temperature ranged from 0 to -5). The Successor Representation's gamma parameter is already theoretically bounded in the range [0, 0.99), and the softmax inverse temperature saturated around 1,200.

Using these parameter boundaries, we simulated behavior from 500 agents for each model (total N = 2,000). Each agent's parameter value(s) were randomly sampled from a uniform distribution, except for the SR inverse temperature, which was sampled from a half-Gaussian distribution by taking the absolute value of samples drawn from a Gaussian distribution with mu = 0 and SD = 400. This procedure allowed us to test whether parameter recovery is acceptable across a broad range (and/or combination) of parameter values, including those that may be uncommon in humans. Using these simulated agents' behaviors, we then estimated parameters using the exact same procedure used to estimate parameters from human subjects' behaviors. For this analysis, we only estimated parameters for a particular model if the agents' behavior was actually generated by that model (e.g., we only estimated parameters for the BFS-backward model for agents whose behavior was simulated from BFS-backward).

Recovery for BFS-backward was adequate (Spearman's $\rho$ = .98), though the optimizer tended to converge upon two (inaccurate) modes when the true search threshold was greater than 10. Recovery was similarly adequate for the BFS-forward model (Spearman's $\rho$ = .97), though the optimizer sometimes overestimated large thresholds (starting at around 12) and underestimates small thresholds (ending at around 4). Recovery for the ideal observer model demonstrated clear biases, such that the optimizer was best at recovering inverse temperatures in the range of [-1, 0], less good at recovering temperatures in the range of [-2, -1], and much worse beyond that. However, the estimates were monotonically related to the true values (Spearman's $\rho$ = .91). The SR's $\gamma$ parameter demonstrated adequate recovery (Spearman's $\rho$ = .81), though smaller values appeared to be a little less reliably recovered, such that true $\gamma < 0.25$ tended to be underestimated as near-zero. Finally, the SR inverse temperature parameter demonstrated mediocre recoverability (Spearman's $\rho$ = .53), such that the optimizer, on occasion, estimated extremely large values of this parameter. Visualizations of all recovery analyses can be found in Figure S3.

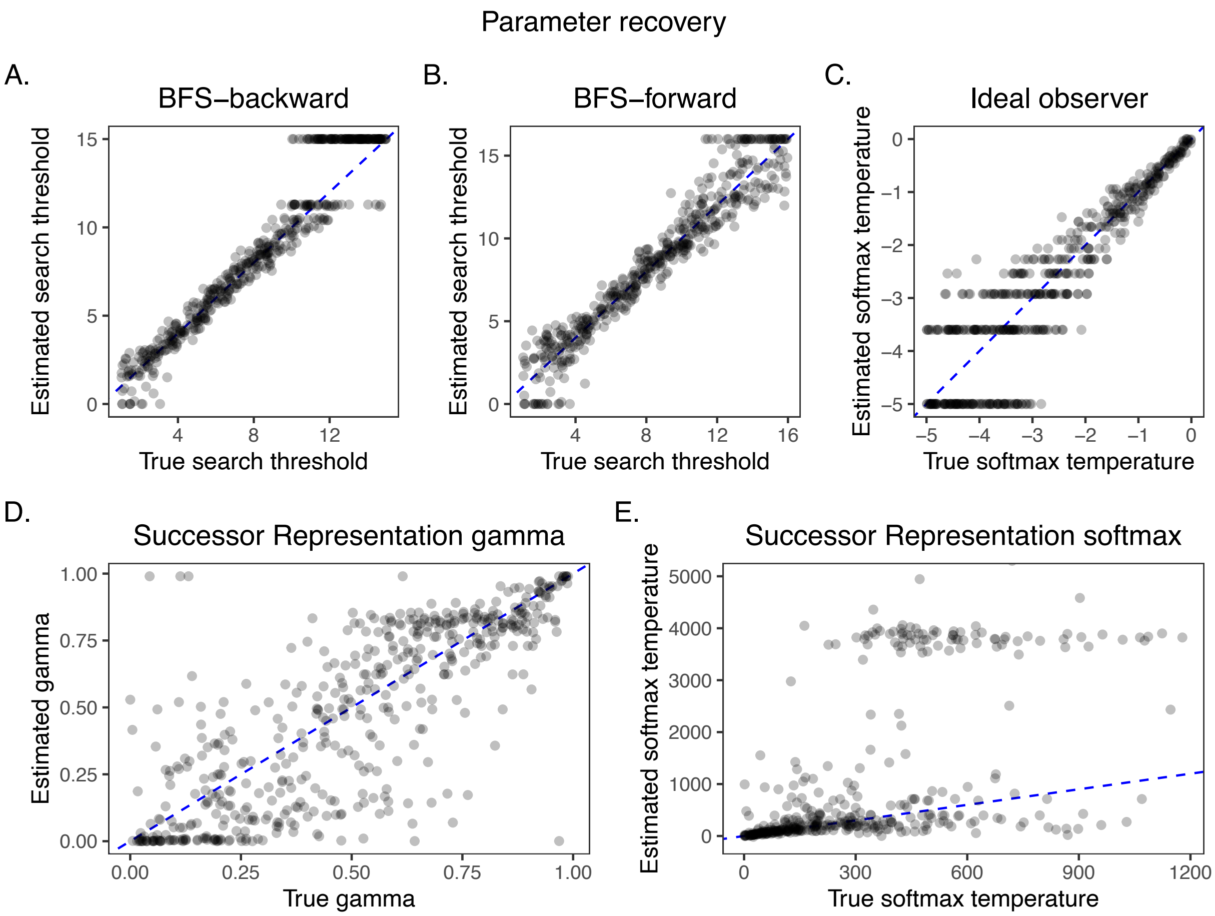


***Figure S3 | Parameter recovery results.*** *The plot of the SR softmax temperature is zoomed in to emphasize more typical values. Dashed blue lines indicate the identity/diagonal line, such that all estimates would fall on this line if parameter recovery were perfect.*

*Estimated parameters.*

Plots of all estimated parameters can be found in Figure S4.

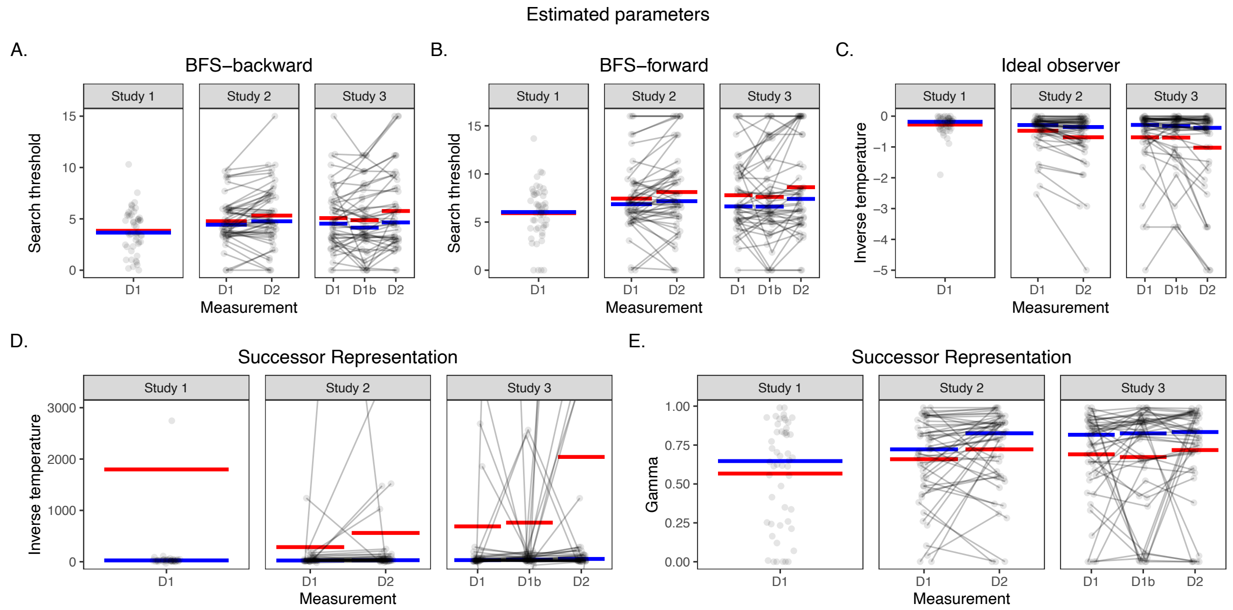


***Figure S4 | Estimated parameters from all computational models.*** *Each individual datapoint reflects one subject, and lines connect subjects' datapoints across repeated measurements. Red crossbars reflect group-level averages, and blue bars reflect medians. To emphasize visualization of more typical values of the SR inverse temperature parameter, the y-axis is truncated to exclude extremely large values. Measurement abbreviations: D1 = before rest on day 1, D1b = after awake rest on day 1, D2 = after overnight rest on day 2.*

*Model confusion.*

To perform the model confusion analysis, we simulated a new dataset of behaviors from agents whose generative parameters were closely matched to real human subjects' estimated parameters, as inferences about model confusability are strongly conditional on the underlying distributions of parameters used to simulate behaviors[1]. For each computational model under consideration, we simulated 500 agents' behaviors. Parameters were sampled (with replacement) from the empirical distribution of estimated parameters from human subjects, and we added a small amount of Gaussian noise to these sampled parameters to increase variability (i.e., because the number of agents outnumbered the number of actual human subjects). Gaussian noise was added to all models with mu = 0, and with SD = 0.5 for the BFS-backward and BFS-forward models' search threshold parameter, SD = 0.1 for the ideal observer model's inverse temperature parameter, SD = 0.04 for the SR's gamma parameter, and SD = 5 for the SR inverse temperature parameter.

After simulating these agents' behaviors (total N = 2,000), we estimated parameters for every model, for every agent, using the exact same procedure used to estimate parameters from human subjects' behaviors. We then classified an agent's behavior as being best-described by a particular model if that model produced the lowest AICc[2] (i.e., Akaike Information Criterion corrected for a small number of observations), which allowed us to construct a confusion matrix quantifying p(best-fitting model = Y | true model = X). We also computed the 'inversion' matrix quantifying p(true model = Y | true model = X), which is useful when interpreting parameter estimates from real subjects' data, as the true model is unknown[1].

The confusion matrix indicates that the SR is confusable with the other models in our set, such that it was only 57% likely to be identified as the best-fitting model when it was actually the true data-generating model (Figure S5), and relatively likely to be misclassified as BFS-backward (14%) or BFS-forward (19%). However, other models are unlikely to be misclassified as the SR (< 10%). This is also reflected in the inversion matrix (Figure S5), such that there is a relatively high chance that the true model is the SR when BFS-backward or -forward is estimated to be the best-fitting model (13% and 17%, respectively), but a relatively small chance of misclassification if the SR is estimated to be the best-fitting model (9-10% per alternative model). Together, these results suggest that the parameter estimation procedure is likely to be biased against the SR.

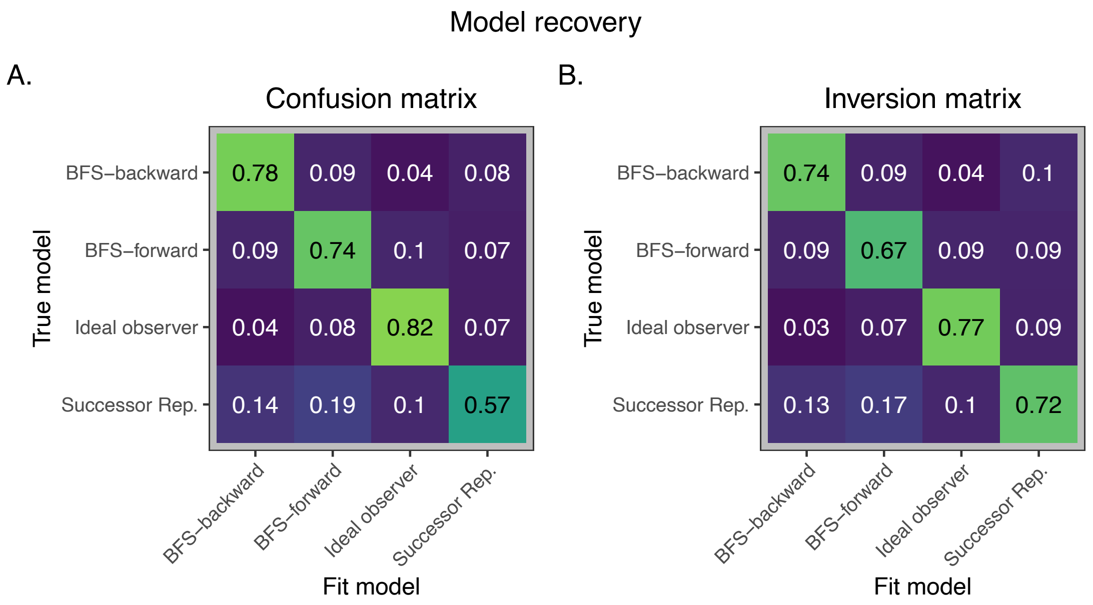


***Figure S5 | Confusion and inversion matrices.*** *When the Successor Representation is the true model, it tends to be confusable with the other models, particularly the two BFS models.*

### MODEL COMPARISON

*Different methods for implementing the SR.*

In principle, the SR could be built using either an analytic or delta-rule method[3]. To test whether one implementation clearly outperformed the other, we fit parameters to both models. Both implementations produced nearly identical likelihoods (Figure S6), and estimated nearly identical values of $\gamma$ (Figure S7). Therefore, our current data cannot distinguish between these possibilities.

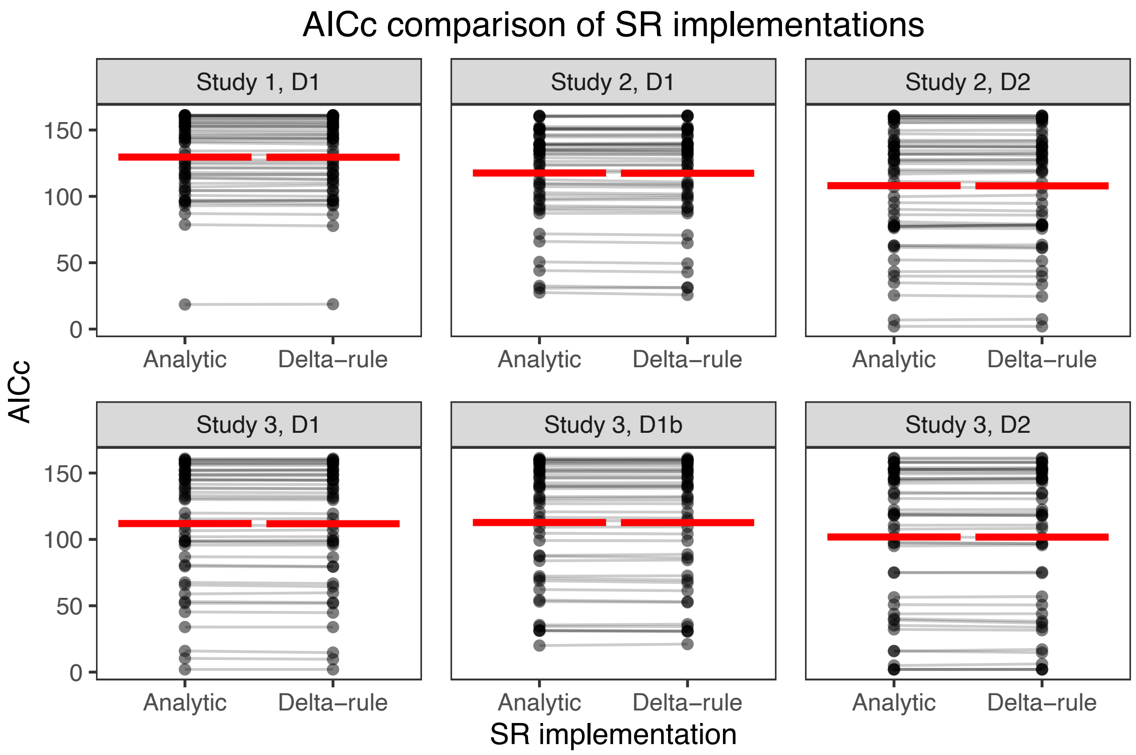


***Figure S6 | Different SR implementations have nearly-identical likelihoods.*** *Each individual datapoint reflects one subject; lines connect each subject's datapoints across the two SR implementations. Red crossbars reflect group-level averages. All lines are nearly parallel, indicating that the likelihoods are functionally identical. AICc stands for Akaike Information Criterion, corrected for a small number of trials. Measurement abbreviations: D1 = before rest on day 1, D1b = after awake rest on day 1, D2 = after overnight rest on day 2.*

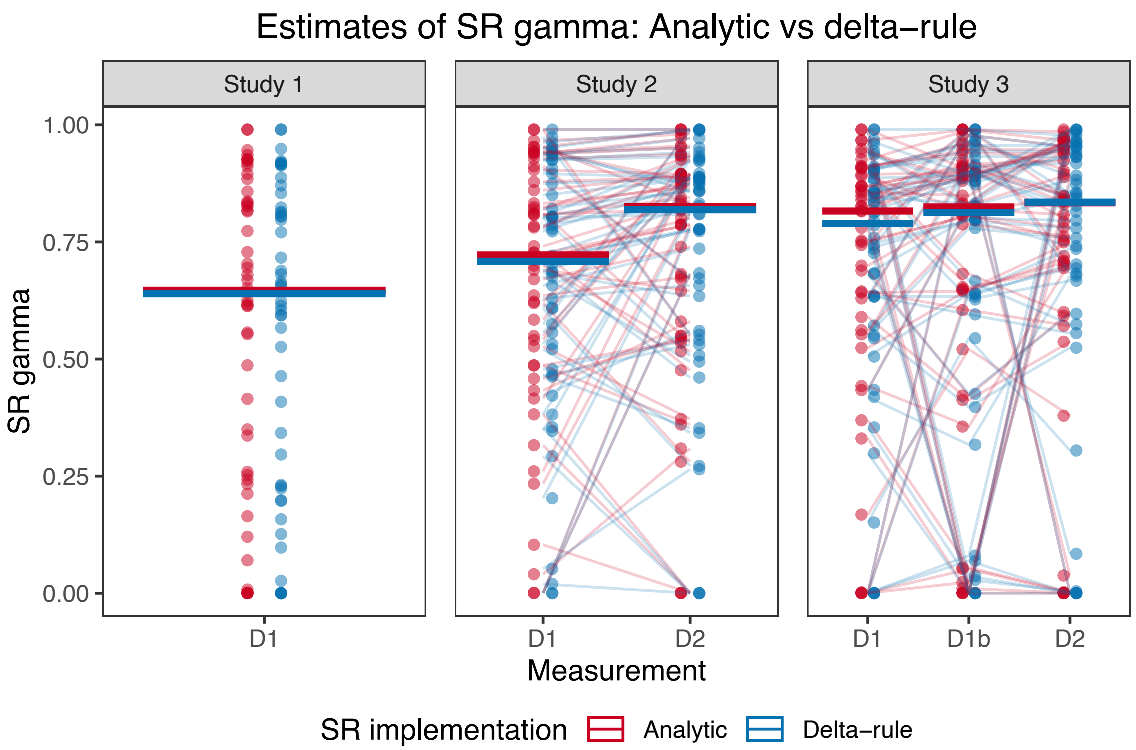


***Figure S7* | *Stable estimates of gamma across two implementations of the SR.*** *Individual datapoints reflect subjects, and lines connect subjects' datapoints in Studies 2-3, where there were repeated measurements. Crossbars reflect group-level medians. Measurement abbreviations: D1 = before rest on day 1, D1b = after awake rest on day 1, D2 = after overnight rest on day 2.*

*McFadden's pseudo-$R^2$.*

In the main text, our model comparison procedure focused on comparing how well the candidate models fit the data, relative to one another. McFadden's pseudo-$R^2$ provides a useful intuition for how well the models fit the data in more absolute terms (Figure S8). The 'null model' used to compute relative gains in $R^2$ was an agent that chose completely at random on each trial.

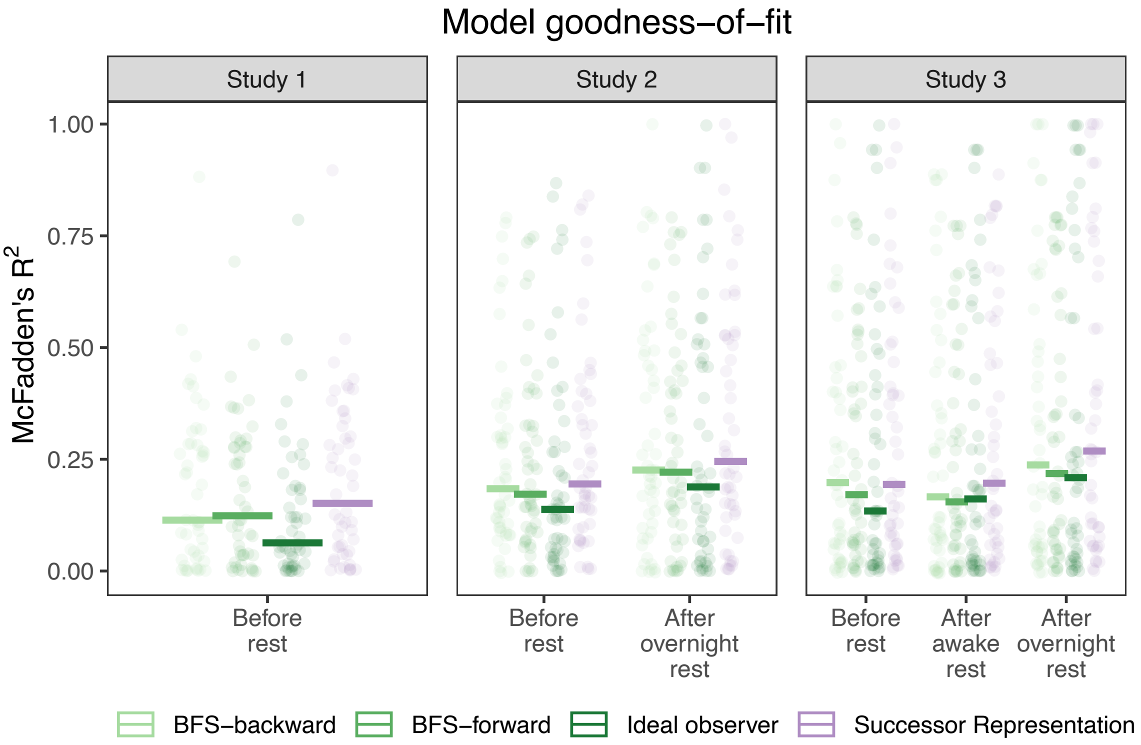


***Figure S8 | McFadden's pseudo $R^2$.*** *Each individual datapoint reflects one subject. Crossbars reflect group-level medians. Gains in $R^2$ are relative to a null model in which the agent chooses completely at random.*

*Disaggregated model comparison metrics.*

In the main text, we report model comparison metrics averaged across studies. Here, we report the same metrics, disaggregated (Figure S9).

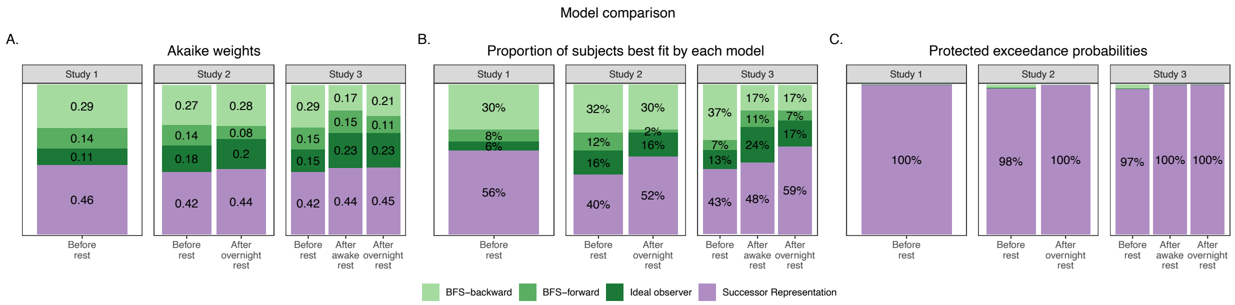


***Figure S9 | Disaggregated model comparison metrics.***

Akaike weights can be interpreted as conditional probabilities for each model[2], which we used to provide evidence for group-level trends. Akaike weights can also be interpreted for individual subjects (Figure S10), which can provide intuitions about subject-level variability.

In the main text, we included a posterior predictive check (PPC) visualization for navigation behaviors before rest. Here, we visualize PPCs for all other timepoints in Figure S11.

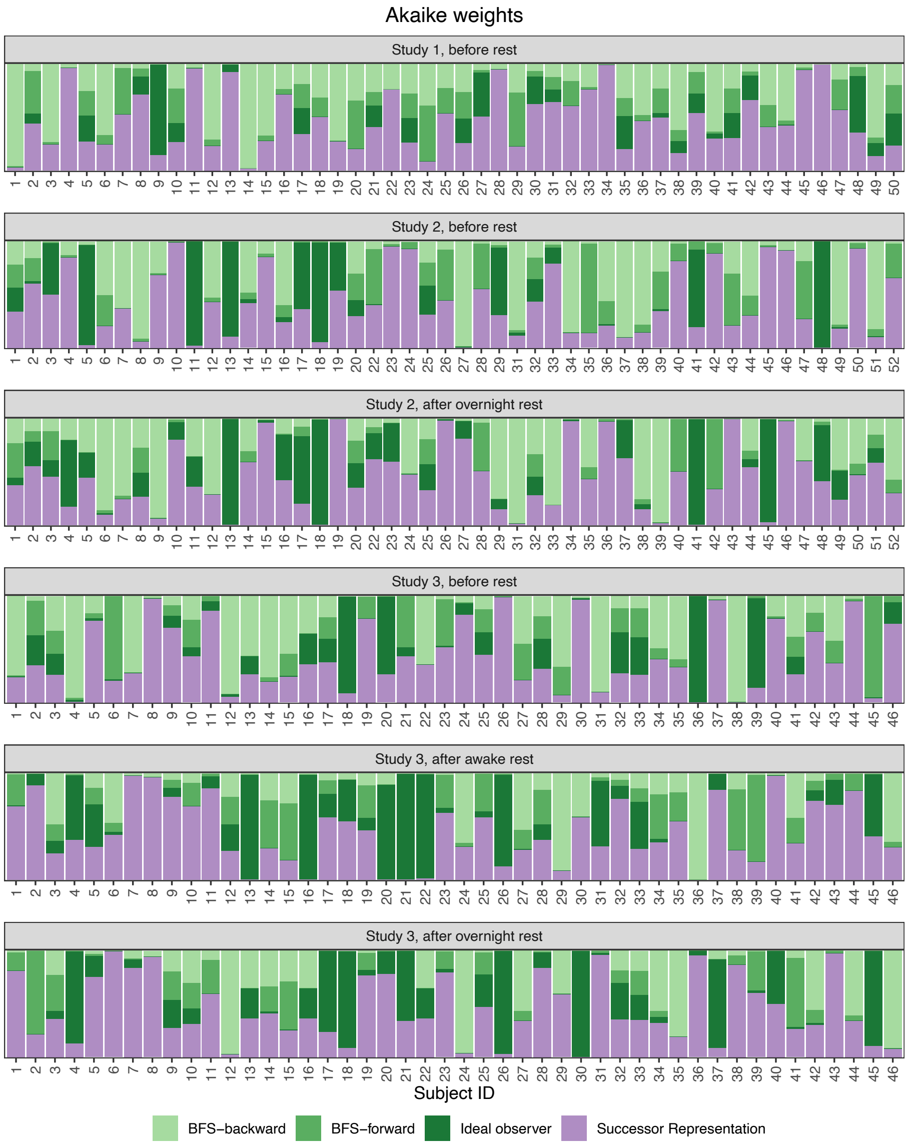


***Figure S10 | Akaike weights per subject, per measurement.***

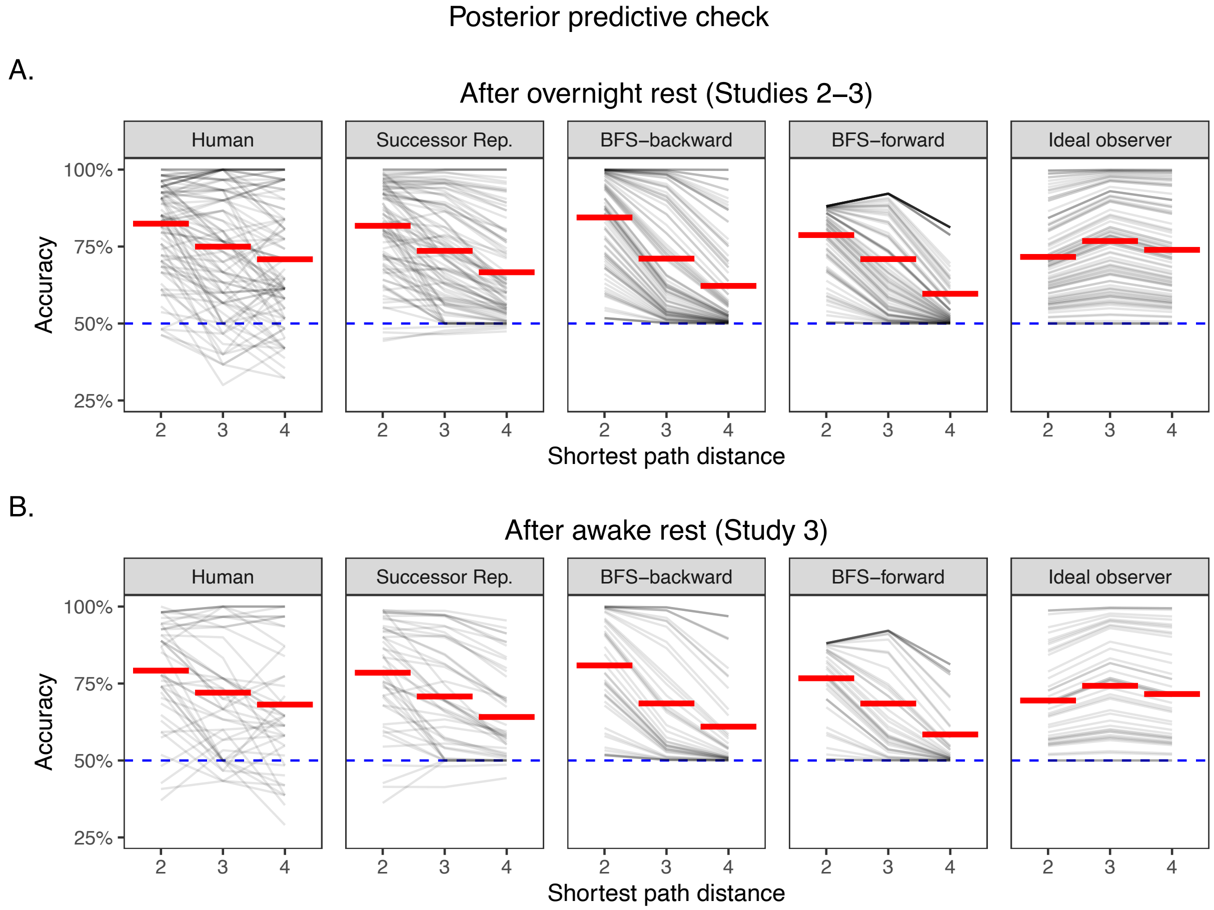


***Figure S11 | Posterior predictive checks for predicted navigation accuracy after overnight and awake rest.***

*Out-of-sample prediction.*

In the main text, we reported out-of-sample likelihoods for a subset of held-out trials in which there were two correct answers (i.e., where both Sources had the same shortest path distance to the Target). Here, we report disaggregated likelihoods in Figure S12.

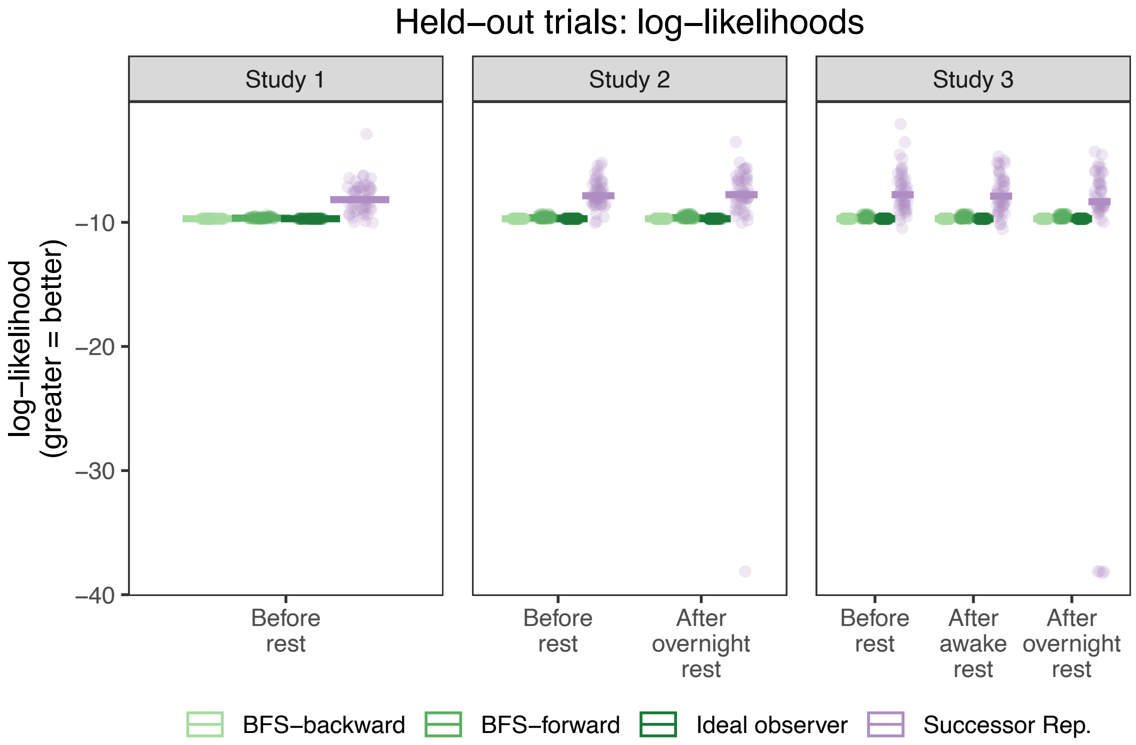


***Figure S12 | The SR best explains human behavior on held-out trials.*** *In a subset of 'held-out' navigation problems (i.e., not used to fit parameters), subjects were presented with two Sources that were the same path distance away from the Target. Despite this, humans frequently demonstrated a preference for one Source over the other. On these held-out trials, the SR has greater out-of-sample likelihoods than the planning models, demonstrating that it is better able to explain human subjects' preferences. Each individual datapoint reflects one subject. Crossbars reflect group-level medians.*

In a follow-up analysis, we tested at which path distances human behavior is consistent with the predictions made by the SR. While the out-of-sample likelihoods demonstrate that the SR is more likely than other models, it is also informative to test how strongly human behavior matches the SR's predictions in terms of absolute accuracy. Results from mixed-effects logistic regression reveal that humans are significantly more likely to choose the SR-preferred Source on longer-range navigation problems (distance-3 $p < .001$, distance-4 $p = .022$; Figure S13), but not distance-2 problems where each of the Sources is directly connected to the Target ($p = .971$; Figure S13).

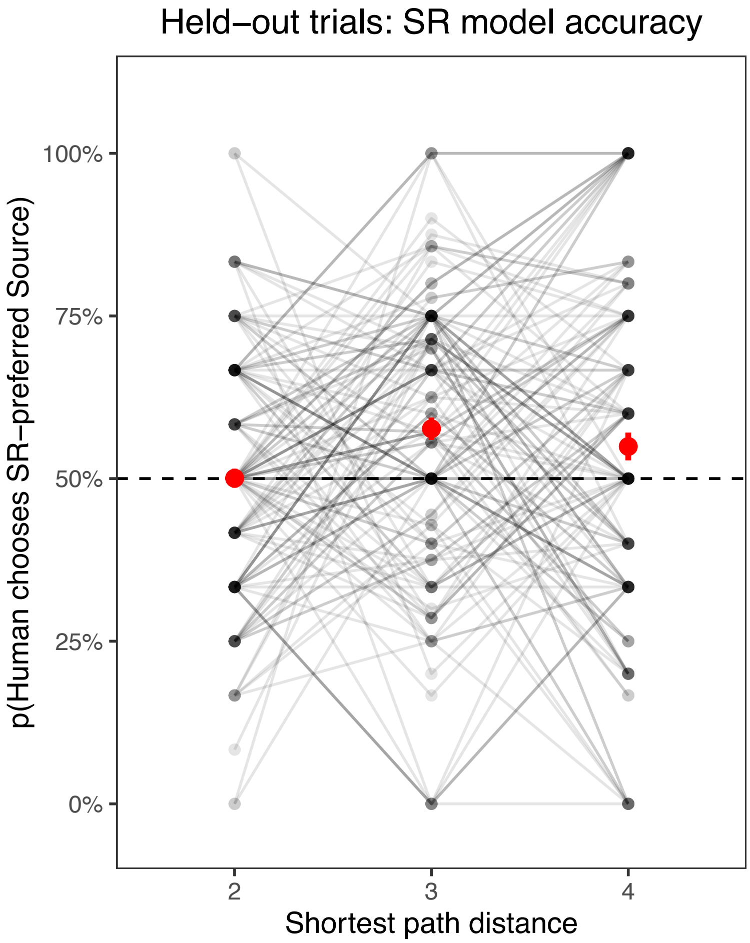


***Figure S13 | Human behavior is consistent with the SR's predictions when there are two 'correct' answers.*** *In a subset of 'held-out' navigation problems (i.e., not used to fit parameters), subjects were presented with two Sources that were the same path distance away from the Target. Although planning models generally predict indifference on these trials, the SR often favors one Source over the other. At the group level, humans demonstrate a slight but significant preference for the SR-preferred Source on longer-range navigation problems, but not the distance-2 problems where both Sources are directly connected to the Target. Each individual datapoint reflects one subject. Red dots reflect the group-level average, estimated from mixed-effects logistic regression; error bars reflect standard error.*

### TRANSITION REEVALUATION

*Social navigation after transition reevaluation.*

The transition reevaluation procedure was administered at the end of Day 2, after subjects had already completed the social navigation task once (Figure 1F). The set of trials used in the post-reevaluation navigation task differed from the set of trials used at all other timepoints, inadvertently introducing the potential confound that the post-reevaluation trials may have simply been more difficult than the main navigation trials (Figure S14). Indeed, simulation results indicate that a fully-asymptotic SR-agent would have worse accuracy in the post-reevaluation trials, despite using the same parameters to solve the pre- and post-reevaluation trials (Figure S14).

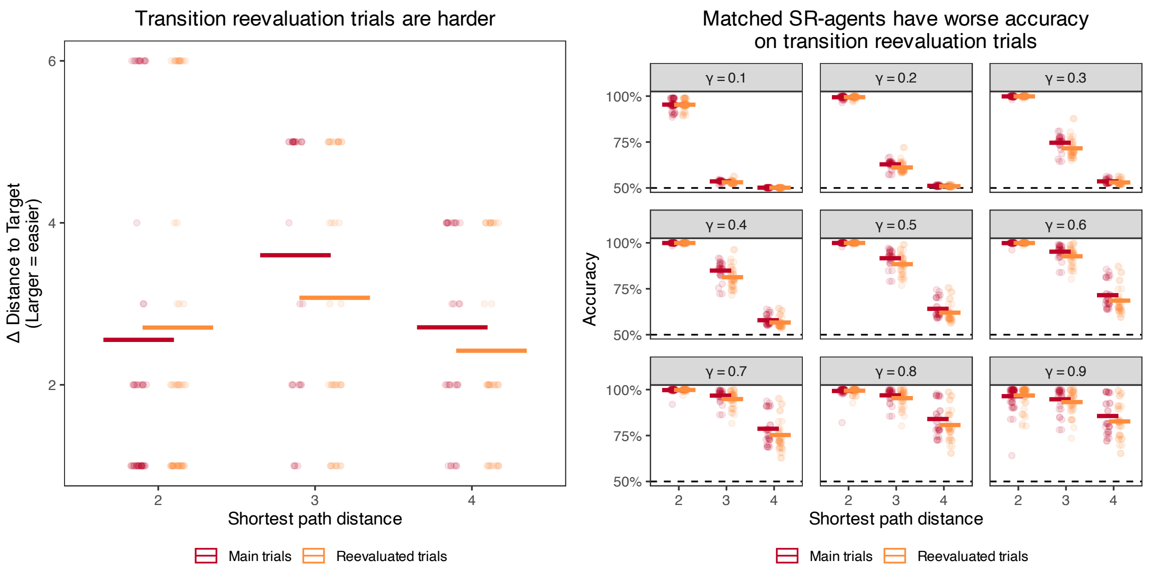


***Figure S14 | Evidence of a task difficulty confound in the transition reevaluation trials. Left:*** *Navigation trials are generally easier to solve when one of the two Sources is very far from the Target. This can be quantified in a 'model-free' way by calculating the absolute difference between the shortest path distances from each Source to the Target (e.g., dist(Source B, Target) – dist(Source A, Target). The larger the difference, the easier the trial. On average, longer-range problems are harder in the post-reevaluation trials (orange) than in the pre-reevaluation trials (red).* ***Right:*** *The same SR-agent tends to performs worse in the post-reevaluation trials.*

Therefore, when testing whether subjects' navigation accuracy decreased following transition reevaluation, we included a trial-by-trial predictor indexing the difference in the two Sources' distances to the Target (Figure S14). We reasoned that this metric provided a reasonable 'model-free' estimate of how difficult each trial was, and that by regressing it out, we could test whether subjects' navigation accuracy decreased following transition reevaluation, above-and-beyond differences in task difficulty. Disaggregated results are reported in Table S6. Of particular note is that following reevaluation, subjects' navigation accuracy was statistically indistinguishable from their accuracy on Day 1 (i.e., before overnight rest), adjusting for differences in task difficulty.

| ***Study*** | ***Reference category*** | ***Term*** | ***Beta estimate*** | ***Z*** | ***95% CI*** | ***p*** |
|---|---|---|---|---|---|---|
| 2 | Distance 2 | Intercept (TR) | 1.70 | 9.36 | [1.35, 2.06] | < .001 |
| | | ΔDistance | 0.05 | 4.69 | [0.03, 0.07] | < .001 |
| | | Distance 3 | -0.67 | -9.19 | [-0.82, -0.53] | < .001 |
| | | Distance 4 | -1.25 | -17.20 | [-1.39, -1.11] | < .001 |
| | | BOR | -0.11 | -0.96 | [-0.34, 0.12] | .338 |
| | | Distance 3 × BOR | 0.04 | 0.40 | [-0.17, 0.26] | .687 |
| | | Distance 4 × BOR | 0.23 | 2.14 | [0.02, 0.43] | .033 |
| | | AOR | 0.10 | 0.89 | [-0.12, 0.31] | .374 |
| | | Distance 3 × AOR | 0.18 | 1.57 | [-0.04, 0.40] | .116 |
| | | Distance 4 × AOR | 0.47 | 4.28 | [0.25, 0.68] | < .001 |
| | Distance 3 | Intercept (TR) | 1.03 | 5.62 | [0.67, 1.39] | < .001 |
| | | Distance 4 | -0.57 | -7.71 | [-0.72, -0.43] | < .001 |
| | | BOR | -0.07 | -0.56 | [-0.31, 0.17] | 0.575 |
| | | Distance 4 × BOR | 0.18 | 1.64 | [-0.04, 0.40] | .102 |
| | | AOR | 0.27 | 2.36 | [0.05, 0.50] | .018 |
| | | Distance 4 × AOR | 0.29 | 2.51 | [0.06, 0.52] | .012 |
| | Distance 4 | Intercept (TR) | 0.46 | 2.52 | [0.10, 0.81] | .012 |
| | | BOR | 0.11 | 0.98 | [-0.12, 0.34] | .328 |
| | | AOR | 0.56 | 5.07 | [0.35, 0.78] | < .001 |
| 3 | Distance 2 | Intercept (TR) | 1.77 | 7.61 | [1.32, 2.23] | < .001 |
| | | ΔDistance | 0.06 | 4.79 | [0.03, 0.08] | < .001 |
| | | Distance 3 | -0.56 | -7.21 | [-0.71, -0.41] | < .001 |
| | | Distance 4 | -0.92 | -11.92 | [-1.08, -0.77] | < .001 |
| | | BOR | -0.09 | -0.66 | [-0.35, 0.18] | .512 |
| | | Distance 3 × BOR | 0.04 | 0.36 | [-0.19, 0.27] | .722 |
| | | Distance 4 × BOR | -0.01 | -0.08 | [-0.23, 0.21] | .937 |
| | | AOR | 0.22 | 1.90 | [-0.01, 0.44] | .058 |
| | | Distance 3 × AOR | -0.10 | -0.88 | [-0.33, 0.13] | .378 |
| | | Distance 4 × AOR | 0.15 | 1.29 | [-0.08, 0.38] | .198 |
| | Distance 3 | Intercept (TR) | 1.21 | 5.17 | [0.75, 1.67] | < .001 |
| | | Distance 4 | -0.36 | -4.48 | [-0.52, -0.20] | < .001 |
| | | BOR | -0.05 | -0.34 | [-0.32, 0.23] | .738 |
| | | Distance 4 × BOR | -0.05 | -0.41 | [-0.29, 0.19] | .679 |
| | | AOR | 0.11 | 0.94 | [-0.12, 0.35] | .349 |
| | | Distance 4 × AOR | 0.25 | 2.05 | [0.01, 0.50] | .040 |
| | Distance 4 | Intercept (TR) | 0.85 | 3.64 | [0.39, 1.30] | < .001 |
| | | BOR | -0.10 | -0.71 | [-0.36, 0.17] | .477 |
| | | AOR | 0.37 | 3.14 | [0.14, 0.59] | .002 |

***Table S6 | Social navigation after reevaluation.*** *Results from mixed-effects logistic regression models including random intercepts and random slopes for the measurement ID (i.e., transition reevaluation, before overnight rest, after overnight rest). 'Transition reevaluation' is abbreviated*

*TR, 'before overnight rest' is abbreviated BOR, and 'after overnight rest' is abbreviated AOR. Note that the reference category was set to TR, such that all 'main effects' reflect estimates for TR, and all 'interaction' effects reflect estimates for BOR or AOR. ΔDistance refers to the absolute difference in the shortest path distances between each Source and the Target, e.g., dist(Source B, Target) – dist(Source A, Target).*

*Simulation of 'on-task' replay.*

In the main text, we use simulation to test whether on-task SR-replay could in principle explain how subjects were able to achieve relatively accurate social navigation following transition reevaluation[4] (Figure 4C). Past studies of the SR have used similar paradigms to investigate nonsocial navigation decisions, and found comparable patterns of human behavior, in which people appear capable of adapting rapidly to changes in their environment[5]. This past work has speculated that following transition reevaluation, humans may be able to quickly engage in enough replay that they can build an updated representation of the multistep relations in the environment. Our simulations suggest that this is a computationally-feasible hypothesis in the context of social navigation, though it is likely that neural data are necessary to definitively address this hypothesis[4].

In the main text, we present results from a simulation study where SR-replay uses a delta-rule mechanism to perform 'on-policy' updates (Equation 1). We note that there are alternative updating rules that could be used for SR-replay, such as using a delta-rule mechanism to estimate the true one-step transition structure of the network, then computing an asymptotic 'off-policy' SR using an analytic, closed-form solution (Equation 2). This kind of off-policy mechanism is not limited to replay-based updating, but could also be used in the course of direct experience with the environment. Simulation results comparing the 'on-policy'/delta-rule SR against the 'off-policy'/analytic SR demonstrate that these different implementations can in principle make diverging predictions (Figure S15). However, given that our parameter-fitting procedure estimated near-identical likelihoods and parameter values for delta-rule and analytic SRs more generally (Figure S6; Figure S7), our current data are unable to adjudicate between these methods.

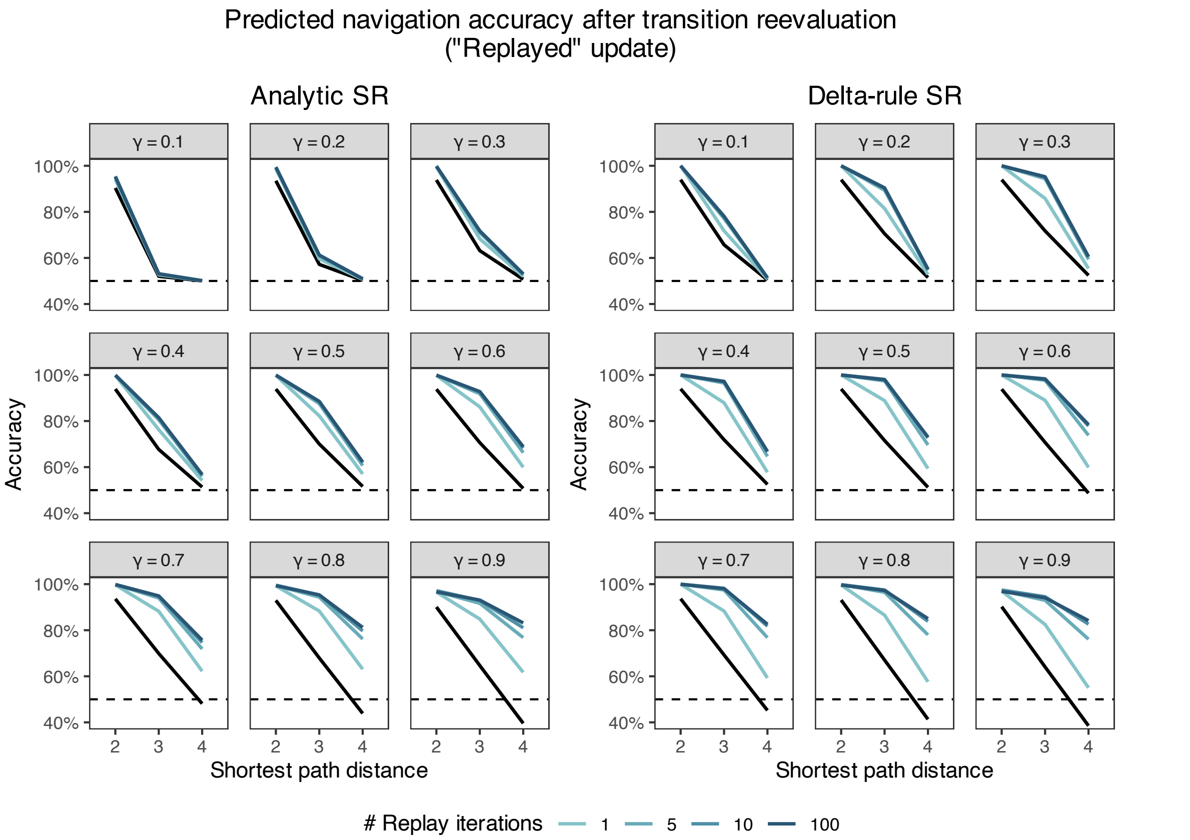


***Figure S15 | Simulated SR-replay from an off-policy/analytic SR-agent, compared to an on-policy/delta-rule SR-agent.*** *Different implementations of the SR make different predictions about behavior after replay due to transition reevaluation, particularly at lower values of* $\gamma$*. The black lines reflect predicted accuracy for an agent that does no updating. The softmax inverse temperature is fixed at* $\beta = 100$ *for visualization.*

Finally, the SR-replay simulation also raises interesting questions about the successor horizon $\gamma$, which dictates the degree of multistep abstraction that is encoded into a representation. If an agent were fully able to control this parameter, it would be able to 'tune' its learning according to what experiences are available to it. For example, when rapidly updating cached representations following a change in the network structure, it intuitively seems that the agent might want to 'propagate' that change over many steps, as to maximize its use of limited directed experience[6]. However, past research suggests that $\gamma$ may (at least in part) function more as a descriptive parameter reflecting how easily an agent can learn multistep abstractions from experience[7], rather than a parameter that an agent can control. This has particularly interesting implications for our finding that $\gamma$ increases after overnight rest. It is possible that during extended periods of rest, such as sleep, increased abstraction occurs from the brain replaying longer random walks through the network. Another (not mutually-exclusive) possibility is that replay of extended experience continues to comprise relatively short sequences[8], but that the temporal proximity of replayed items results in greater overlap of neural patterns associated with each item[6,9-11], and therefore more optimal conditions for encoding multistep relations.

*Simulation of surprise-driven 'single-shot' re-caching.*

We note that SR-replay is not the only mechanism through which a subject's representation could be re-cached following transition reevaluation. For example, given that it is surprising to observe sudden changes in network members' friendships, it is possible that a 'single-shot' observation of these changes could be encoded with a higher learning rate (i.e., greater $\alpha$ in the delta-rule implementation of the SR commonly used in reinforcement learning)[12]. This mechanism is not mutually exclusive with replay; indeed, a greater learning rate would be a natural complement to surprise-evoked replay. To test how much a 'single-shot' learning update could help an agent to re-cache its representation, we simulated the effect of observing nodes 7 and 13 becoming friends, and nodes 1 and 2 breaking off their friendship. As the SR's core updating mechanism is observing co-occurrences, rather than the *absence* of co-occurrence, we modeled the latter as fictive observations (or replay) of all of node 1 and 2's friendships (i.e., $[1 \rightarrow 8]$, $[2 \rightarrow 3]$, $[2 \rightarrow 4]$), excluding the friendship that had just been broken (i.e., $[1 \rightarrow 2]$).

Simulation results confirm that SR-agents that do any amount of updating demonstrate some amount of improvement on the longer-distance problems (i.e., distance-3 and -4), aside from agents with the lowest values of gamma (Figure S16). In the best-case scenario, a single-shot update can produce surprisingly large gains in longer-distance accuracy. While these agents are far from achieving ceiling-level accuracy, these results demonstrate that an SR-agent can achieve above-chance navigation accuracy simply by encoding what was instructed during the transition reevaluation procedure. Therefore, single-shot updating is sufficient for explaining how humans might achieve above-chance accuracy after transition reevaluation.

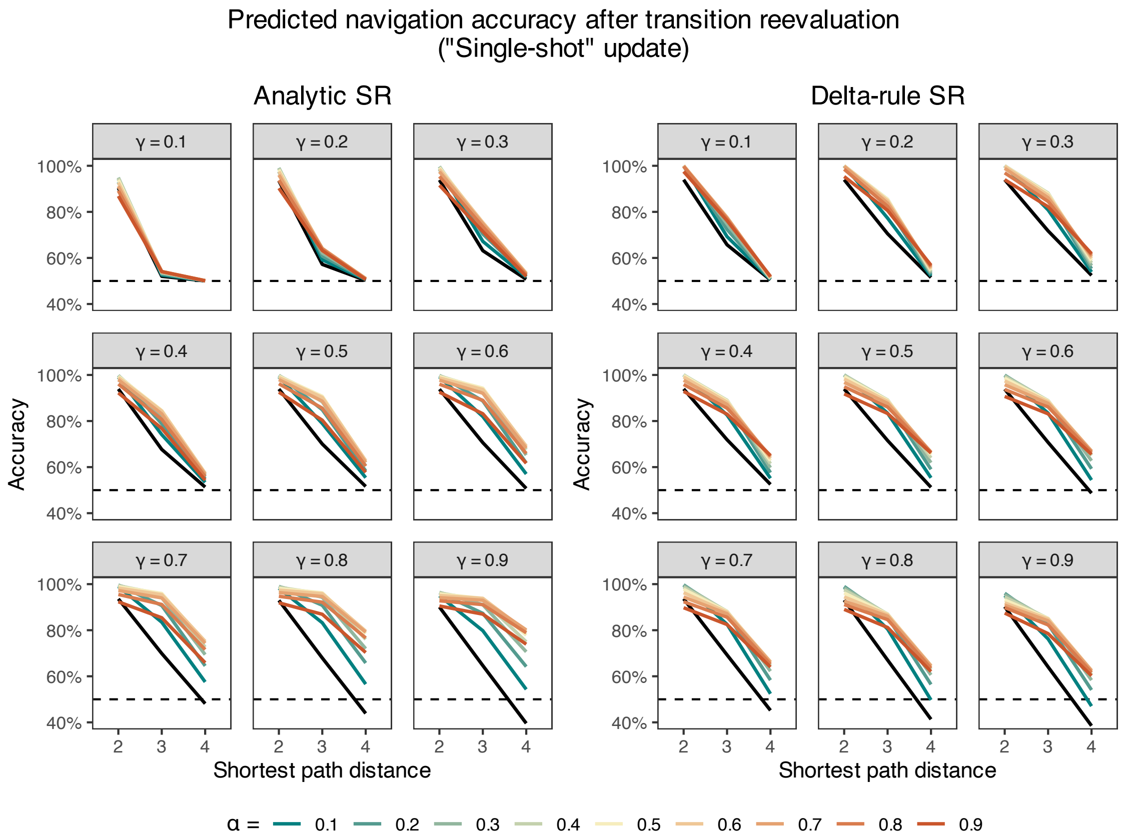


***Figure S16 | Simulated 'single-shot' updating.*** *Whether the SR-agent uses an off-policy/analytic method, or an on-policy/delta-rule method, a single-shot update is sufficient for explaining how people might achieve above-chance accuracy on longer-distance navigation problems. The black lines reflect predicted accuracy for an agent that does no updating. Generally, higher learning rates (α) are associated with greater accuracy. The softmax inverse temperature is fixed at* $\beta = 100$ *for visualization.*

### SOCIAL PRIORS IN LEARNING

In our study, we explicitly instructed subjects that they were learning about a social network. Our hypothesis was that people use the domain-general mechanism of multistep abstraction to learn abstract cognitive maps of social networks, which can then be used to make social decisions. Therefore, if subjects were to demonstrate similar behaviors in (e.g.) a task where they learned about a network of computers passing messages to each other, this would not contradict our hypothesis. However, it would be far more problematic for our hypothesis if subjects did not construe the task as having any social element whatsoever. We reasoned that there may be uniquely social prior beliefs that might bias how people learn about friendships in a social network. If true, this would suggest that subjects were using a domain-general mechanism to navigate through what they perceived as being a specifically *social* network.

The learning task required subjects to actively guess (or report) whether a Target was friends with all of the other network members. We therefore analyzed subjects' responses to all non-friends in order to test whether subjects were more likely to guess (nonexistent) friendships between network members that were the same gender and/or race. Although the figures in our manuscript use cartoon avatars, we note that the actual stimuli used in our study were photographs of real humans drawn from a racially-diverse stimulus set[13] and given demographically-appropriate names, enhancing the social realism of the task.

The learning task consisted of six blocks in total, where each block presented every network member as a Target. On each trial, subjects were presented with virtual 'flash cards'. When shown a Target network member, participants were required to find all of the Target's friends from the face-down cards. When a card corresponding to the Target's friend was selected, the card flipped face-up to reveal their photograph. Cards remained face-down in response to incorrect guesses. To ensure that subjects were truly learning about friendships, rather than something low-level like the cards' spatial location, we shuffled where the cards were located in block four. Due to this task design, subjects were only able to guess randomly during blocks one and four, and we therefore excluded these blocks from our analysis.

Results from mixed-effects logistic regression reveal that learning was robustly biased by social priors: subjects were significantly more likely to incorrectly guess friendship between network members if they were the same gender ($b = 0.17$, $Z = 5.70$, 95% CI = [0.11, 0.23], $p < .001$), or if they were the same race ($b = 0.17$, $Z = 6.87$, 95% CI = [0.12, 0.22], $p < .001$). Moreover, these priors proved sticky, as we continued to find evidence of these biases even when restricting our analysis to the very last block (gender $b = 0.15$, $Z = 3.77$, 95% CI = [0.07, 0.23], $p < .001$; race $b = 0.11$, $Z = 2.29$, 95% CI = [0.02, 0.21], $p = .022$). These results suggest that subjects' learning about a social network diverged from how they might learn about other kinds of networks, and provide evidence that our task serves as an ecologically-valid testbed of the fundamental computational problems posed by navigation within social networks.

### SUPPLEMENTARY REFERENCES


1 Wilson, R. C. & Collins, A. G. E. Ten simple rules for the computational modeling of behavioral data. *eLife* **8**, e49547 (2019). https://doi.org:10.7554/eLife.49547

2 Wagenmakers, E.-J. & Farrell, S. AIC model selection using Akaike weights. *Psychonomic Bulletin & Review* **11**, 192-196 (2004). https://doi.org:10.3758/BF03206482

3 Momennejad, I. Learning Structures: Predictive Representations, Replay, and Generalization. *Current Opinion in Behavioral Sciences* **32**, 155-166 (2020). https://doi.org:https://doi.org/10.1016/j.cobeha.2020.02.017

4 Wittkuhn, L., Krippner, L. M. & Schuck, N., W. Statistical learning of successor representations is related to on-task replay. *bioRxiv*, 2022.2002.2002.478787 (2022). https://doi.org:10.1101/2022.02.02.478787

5 Momennejad, I. *et al.* The successor representation in human reinforcement learning. *Nature Human Behaviour* **1**, 680-692 (2017). https://doi.org:10.1038/s41562-017-0180-8

6 Gershman, S. J., Moore, C. D., Todd, M. T., Norman, K. A. & Sederberg, P. B. The Successor Representation and Temporal Context. *Neural Computation* **24**, 1553-1568 (2012). https://doi.org:10.1162/NECO_a_00282

7 Son, J.-Y., Bhandari, A. & FeldmanHall, O. Abstract cognitive maps of social network structure aid adaptive inference. *Proceedings of the National Academy of Sciences* **120**, e2310801120 (2023). https://doi.org:10.1073/pnas.2310801120

8 Davidson, T. J., Kloosterman, F. & Wilson, M. A. Hippocampal Replay of Extended Experience. *Neuron* **63**, 497-507 (2009). https://doi.org:10.1016/j.neuron.2009.07.027

9 Lewis, P. A. & Durrant, S. J. Overlapping memory replay during sleep builds cognitive schemata. *Trends in Cognitive Sciences* **15**, 343-351 (2011). https://doi.org:10.1016/j.tics.2011.06.004

10 Kumaran, D. & McClelland, J. L. Generalization through the recurrent interaction of episodic memories: A model of the hippocampal system. *Psychological Review* **119**, 573-616 (2012). https://doi.org:10.1037/a0028681

11 Schapiro, A. C., Rogers, T. T., Cordova, N. I., Turk-Browne, N. B. & Botvinick, M. M. Neural representations of events arise from temporal community structure. *Nature Neuroscience* **16**, 486-492 (2013). https://doi.org:10.1038/nn.3331

12 Lamba, A., Frank, M. J. & FeldmanHall, O. Anxiety Impedes Adaptive Social Learning Under Uncertainty. *Psychological Science* **31**, 592-603 (2020). https://doi.org:10.1177/0956797620910993

13 Ma, D. S., Correll, J. & Wittenbrink, B. The Chicago face database: A free stimulus set of faces and norming data. *Behavior Research Methods* **47**, 1122-1135 (2015). https://doi.org:10.3758/s13428-014-0532-5
